# Maast: genotyping thousands of microbial strains efficiently

**DOI:** 10.1101/2022.07.06.499075

**Authors:** Zhou Jason Shi, Stephen Nayfach, Katherine S. Pollard

**Affiliations:** Chan Zuckerberg Biohub, San Francisco, CA; Gladstone Institutes, Data Science and Biotechnology, San Francisco, CA; Department of Energy, Joint Genome Institute, Walnut Creek, CA; Lawrence Berkeley National Laboratory, Environmental Genomics and Systems Biology Division, Berkeley, CA; University of California San Francisco, Department of Epidemiology and Biostatistics, San Francisco, CA

## Abstract

Genotyping single nucleotide polymorphisms (SNPs) of intraspecific genomes is a prerequisite to performing population genetic analysis and microbial epidemiology. However, existing algorithms fail to scale for species with thousands of sequenced strains, nor do they account for the biased sampling of strains that has produced considerable redundancy in genome databases. Here we present Maast, a tool that reduces the computational burden of SNP genotyping by leveraging this genomic redundancy. Maast implements a novel algorithm to dynamically identify a minimum set of phylogenetically diverse conspecific genomes that contains the maximum number of SNPs above a user-specified allele frequency. Then it uses these genomes to construct a SNP panel for each species. A species’ SNP panel enables Maast to rapidly genotype thousands of strains using a hybrid of whole-genome alignment and k-mer exact matching. Maast works with both genome assemblies and unassembled sequencing reads. Compared to existing genotyping methods, Maast is more accurate and up to two orders of magnitude faster. We demonstrate Maast’s utility on species with thousands of genomes by reconstructing the genetic structure of *Helicobacter pylori* across the globe and tracking *SARS-CoV-2* diversification during the COVID-19 outbreak. Maast is a fast, reliable SNP genotyping tool that empowers population genetic meta-analysis of microbes at an unrivaled scale.

**Availability:** source code of Maast is available at https://github.com/zjshi/Maast.

## Background

Many bacterial and viral species now have thousands of sequenced genomes in public databases, and these numbers are rapidly increasing, fueled by technologies such as metagenome assembly, high-throughput culturing, and single-cell genome sequencing. For many species, genome collections harbor immense genetic variation [1,2]. Single nucleotide polymorphisms (SNPs) are genomic positions that vary between genomes of the same species with a minimum minor allele frequency (e.g., 1%). Vertically inherited SNPs in conserved genes are commonly used to reconstruct phylogenies and study biogeography [3,4], while SNPs in pathogenicity loci and antibiotic resistance genes are leveraged for surveillance of medically important strains [5–7]. Compared to multilocus sequence typing methods, whole-genome SNP genotyping generally enables greater phylogenetic resolution [8]. Comparing genomes based on a pre-defined set of SNPs is also more computationally scalable than using whole-genome nucleotide identity [9] and thus especially well-suited for the analysis of large genome collections.

SNPs are often identified from whole genome sequences. For example, Parsnp [10] constructs multiple sequence alignments of high-quality genome assemblies and identifies variable positions directly from the alignments. An alternative method is to call SNPs from sequencing reads without genome assembly. For example, Snippy (https://github.com/tseemann/snippy) identifies SNPs from the alignment of short reads to a reference genome and kSNP [11] identifies SNPs using informative k-mers found on short reads, contigs, or genome assemblies. Assembly-free genotyping and k-mer matching are usually faster but less accurate than genotyping SNPs from whole-genome alignments [12]. With all of these strategies, SNPs of interest can be extracted using thresholds on the minor allele frequency (MAF), site prevalence, or protein coding change across a set of strains.

Despite the variety of SNP genotyping methods, a rapid increase in the number of sequenced microbial genomes presents a computational challenge for existing tools. Sequence alignment is the major obstacle to analyzing so many genomes, though kSNP also remain largely untested with thousands of strains. A second challenge is the fact that many species have a high level of genome redundancy[13], especially when a biased sample of clonally related genomes has been sequenced, which is common for clinically important pathogens that are under intensive surveillance (e.g., PulseNet [14] and NCBI Pathogen Detection). This redundancy masks the diversity of unevenly sampled species, and it means that strains from poorly sampled lineages contribute little to the discovery of SNPs, especially when a relatively high MAF threshold is used. Redundancy also alters the meaning of MAF and site prevalence; frequency in the sampled genomes is not a good estimate of frequency in the population. Together these two challenges limit the utility of SNP genotyping methods for many microbial species.

To address this gap, we present Maast (Microbial agile accurate SNP Typer) for accurate genotyping of orders of magnitude more microbial strains than other state-of-the-art methods. Our key innovation is an algorithm to pick a minimal set of maximally diverse genomes. Only these genomes are used for SNP discovery which reduces genomic redundancy and computational cost. We also implement a hybrid method combining whole-genome alignment and optimized k-mer exact match for genotyping SNPs in either assembled genomes or unassembled whole-genome sequencing (WGS) libraries. Maast performs SNP genotyping faster and more accurately than existing methods but sometimes misses rare variants present in genomes not selected for SNP discovery. We apply Maast to a large collection of previously sequenced *Helicobacter pylori* (*H. pylori*) strains and summarize the biogeographic patterns of this species across the globe. We also show that Maast can efficiently track the evolution of *SARS-CoV-2* during the COVID-19 outbreak. Maast is available as open-source software with both source code and documentation freely accessible on GitHub (https://github.com/zjshi/Maast).

## Results

### The Maast SNP genotyping pipeline

Maast is an open-source bioinformatics pipeline written in Python and C++ that fully automates SNP calling and genotyping for microbial species. Maast has two components: (1) constructing a reference panel of SNPs for a microbial species using a reduced set of non-redundant genomes (Figure 1a and b), and (2) ultra-fast, *in silico* genotyping of reference SNPs from large scale genome collections. Genotyping can be performed using draft genome assemblies or unassembled short or long reads (Figure 1c,d and S2) and can be applied to any microbial species, such as bacteria, archaea, viruses, or microbial eukaryotes.

**Figure 1.**
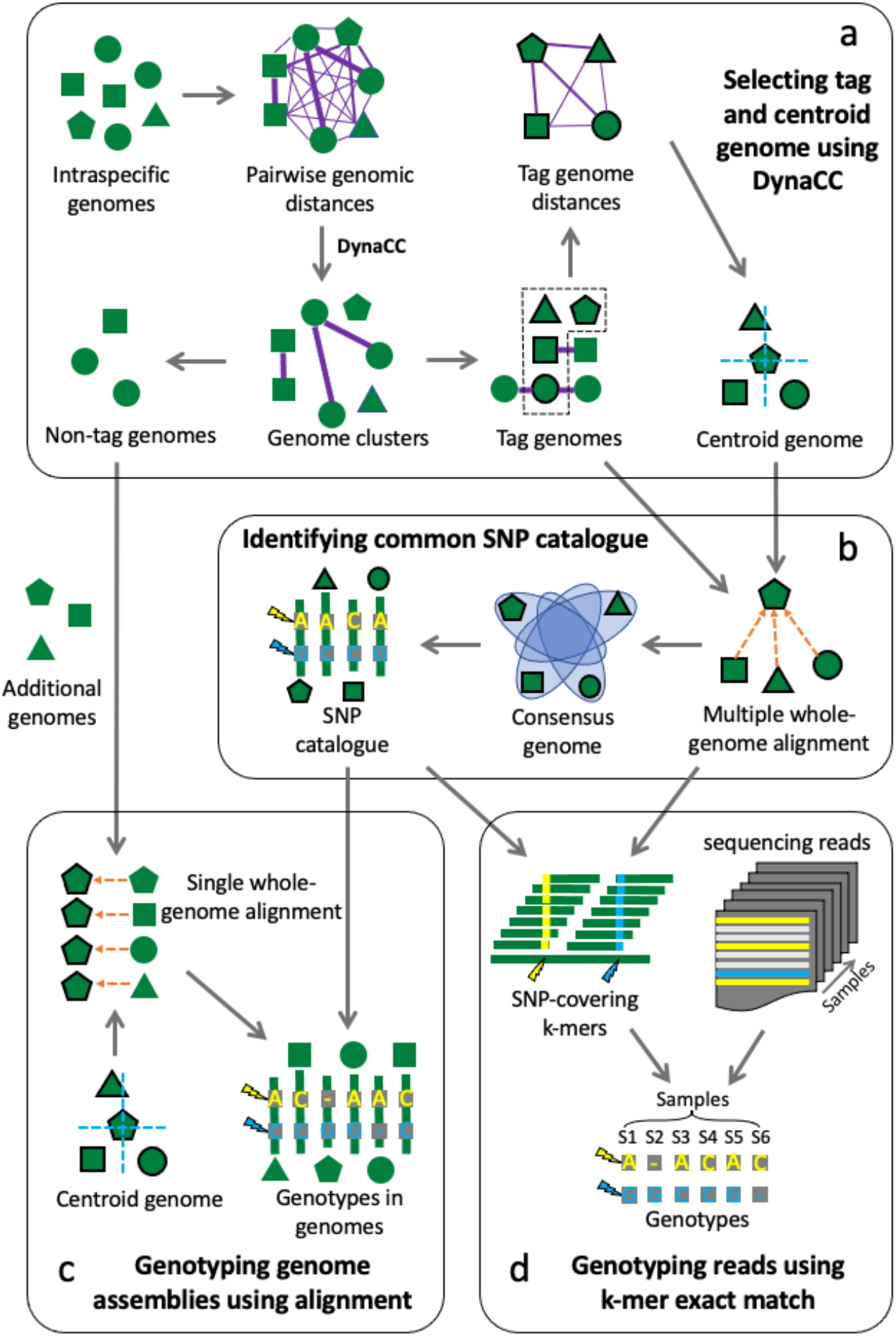
Schematic design of Maast. Maast consists of four major components. (a) Tag genomes and centroid genomes are selected based on Mash distances. A novel algorithm, called DynaCC, is used to automatically choose the genome clustering threshold based on each species’ level of genomic redundancy (see methods). (b) A panel of common SNPs is constructed for each species using multiple whole-genome alignment. (c) SNPs in non-tag genomes or other input genomes are genotyped using single whole-genome alignment. (d) SNPs in short reads are genotyped using k-mer exact matching.

In the first step of the pipeline, Maast rapidly builds a reference SNP panel from a diverse, non-redundant subset of input genome assemblies using MUMmer. The genome subset is identified using pairwise genetic distances and a dynamic graph algorithm that we developed and called DynaCC (Dynamic Connected Component search; Figure S3). DynaCC seeks to identify the minimal number of distinct genomes (i.e., tag genomes) that maximize the number of identified SNPs. From the tag genomes, Maast selects one centroid tag genome to be used as a species reference. Maast then aligns all other tag genomes to the reference using MUMmer, constructs a multiple sequence alignment, and uses a standard SNP calling workflow (Methods) to generate the SNP panel at the user-specified MAF and prevalence (% of genomes containing either SNP allele).

We assessed whether the Maast strategy of subsampling reduces our power to detect SNPs on a benchmark dataset of 146 common bacterial species from the human gut, each with at least 200 high-quality genomes (Table S1). These species have different levels of intraspecific diversity (Figure S4), genomic redundancy (Figure S5a), and SNP density (Figure S5b). Overall, we found that the number of SNPs identified with Maast (>1% MAF) was equal to or greater than the number of SNPs identified using the full set of genomes (Figure 2a,b), with high overlap between the approaches (median=87.9%, Figure 2c). Our results demonstrate that more genomes do not necessarily lead to the discovery of more common SNPs (>1% MAF). The presence of many highly related genomes also leads to biased estimation of SNP frequencies and may reduce SNPs discovered at a given MAF threshold (Figure 2d, e, S6). These analyses highlight that the Maast strategy of subsampling captures most common genetic variation and successfully represents genomes from poorly sampled lineages.

**Figure 2.**
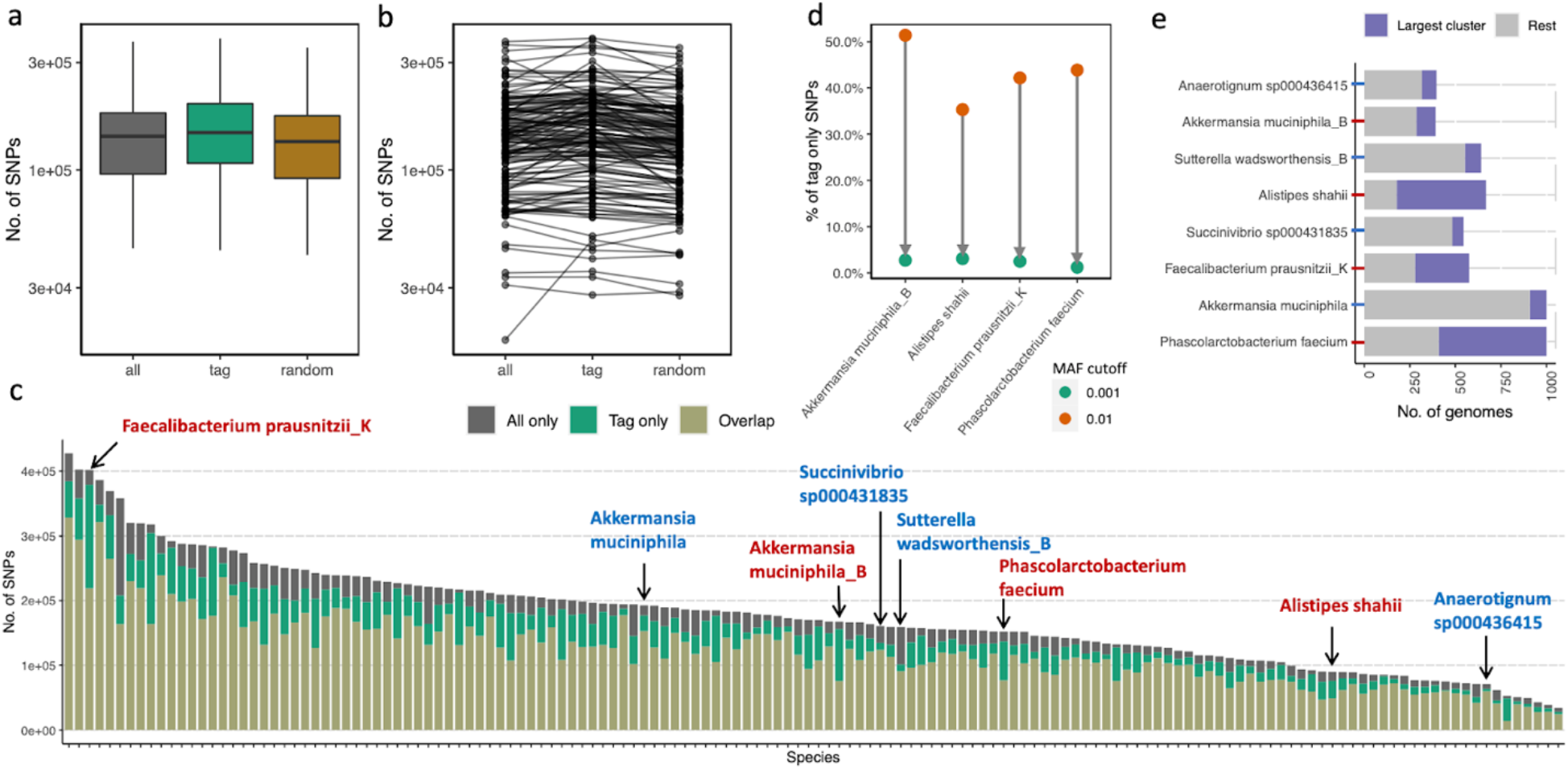
SNP genotyping of 146 human gut bacterial species using tag genomes. (a and b) SNP discovery comparison of Maast with all genomes (grey), only tag genomes (green), and a random set of genomes equal in number to the tag genomes (brown), shows that more genomes do not lead to the discovery of more SNPs. Each box in (a) summarizes the number of SNPs across 146 species. Each point in (b) represents a species, with black lines connecting the data for the same species. For computational efficiency, only the 1,000 highest quality genomes were included for species with >1,000 genomes. (c) Comparison of SNPs discovered by Maast with all genomes versus only tag genomes. Each bar represents a species, with the height of a bar showing the number of SNPs discovered exclusively with all genomes (grey), exclusively with tag genomes (green), or by both approaches (beige). Arrows point to eight example species (from left to right): *Faecalibacterium prausnitzii_K* (species id: 101300), *Akkermansia muciniphila* (102454), *Akkermansia muciniphila_B* (102453), *Succinivibrio sp000431835* (100412), *Sutterella wadsworthensis_B* (101361), *Phascolarctobacterium faecium* (103439), *Alistipes shahii* (100003) and *Anaerotignum sp000436415* (100177). Species label color indicates whether this species has a high (red) or low (blue) level of tag-only SNPs, which is estimated as a fraction of all SNPs that are discovered with tag genomes and not with all genomes. (d and e) SNP sites missing from a small percentage of genome assemblies as they fall below the user-specified prevalence threshold due to being absent in a group of redundant genomes. (d) Examples of species in (c) with a high proportion of tag-only SNPs. The proportion of tag-only SNPs drops if the MAF cutoff for calling SNPs with all genomes is lowered from 0.01 (orange) to 0.001 (green). Most of the SNPs only discovered with tag genomes (tag-only SNPs) are due to MAFs below the 1% threshold in all genomes. (e) The single largest genome cluster is larger for species with many tag-only SNPs compared to those with fewer. The largest genome clusters were compared between species from (c) with high (red axis tick) and low (blue axis tick) levels of tag-only SNPs. For three of four species with more SNPs discovered in the tag-only analysis, the largest cluster contains more than half of all genomes, implying a high level of genome redundancy that biases MAF estimation and leads to an undercount of SNPs. Height of bars shows the total number of genomes in a species and the proportion colored in purple indicates the size of the single largest genome cluster of that species. Every two adjacent species have a similar total number of genomes.

In the second step of the pipeline, Maast performs reference based *in silico* genotyping for new genomes or other genomes in the collection. The workflow can be executed using genome assemblies or unassembled WGS reads. For genome assemblies, Maast aligns each query genome to the centroid genome for that species using MUMmer and directly genotypes alleles for each of the reference SNPs. For WGS reads, Maast performs exact matching between k-mers extracted from the sequencing reads and a database of k-mers covering each SNP in the reference panel (Figure S2). The algorithm is an extension of our previous method GT-Pro [15] for genotyping SNPs in metagenomic sequencing libraries, specifically optimized for WGS of microbial isolates.

### Maast is orders of magnitude faster than existing tools

We evaluated the computational performance of Maast with kSNP3 and Parsnp, two other commonly used methods for microbial SNP calling. All methods were run for a single bacterial species, *Agathobacter rectalis* (Table S2), using 500, 1,000 or 5,000 randomly sampled genome sequences (Methods). Surprisingly, both kSNP3 and Parsnp failed to run through all the *A. rectalis* genomes. kSNP3 was manually terminated after running for a maximum allowed time (48 hours) and consuming more than 1TB disk space, while Parsnp failed to run even for 1,000 genomes, apparently due to high intraspecific genetic diversity. To check that these results were not specific to *A. rectalis*, we ran both tools on two other human gut species (Table S2): *Alistipes putredinis* (n=3,646) and *Bacteroides_B dorei* (n=3,170) and obtained the same result. We conclude that, to our knowledge, Maast is the only tool able to call SNPs in species with thousands of diverse genomes.

To evaluate computational performance when genomic divergence is more limited and in the absence of structural variation, we simulated 5,000 whole genome sequences of *A. rectalis* by randomly introducing SNPs across a representative genome (Methods). While Parsnp ran to completion, kSNP3 again exceeded our 48-hour maximum time window. Overall, Maast was 6.3 to >127 fold faster than Parsnp and 31 to >127 fold faster than kSNP3 (Fig. 3a). In addition, Maast required only 7.2 GB of RAM to process the 5,000 genomes, which is substantially less than what Parsnp used to process 100 genomes and similar to what kSNP3 used to process 1,000 genomes (Table S3). Maast also only used a moderate amount (~1.2GB) of disk space. We attribute Maast’s high speed and RAM-efficiency mainly to the strategy of subsampling genomes for SNP discovery. Other factors included compact data structures and parallel processing. These results demonstrate that Maast can easily run on a personal computer.

**Figure 3.**
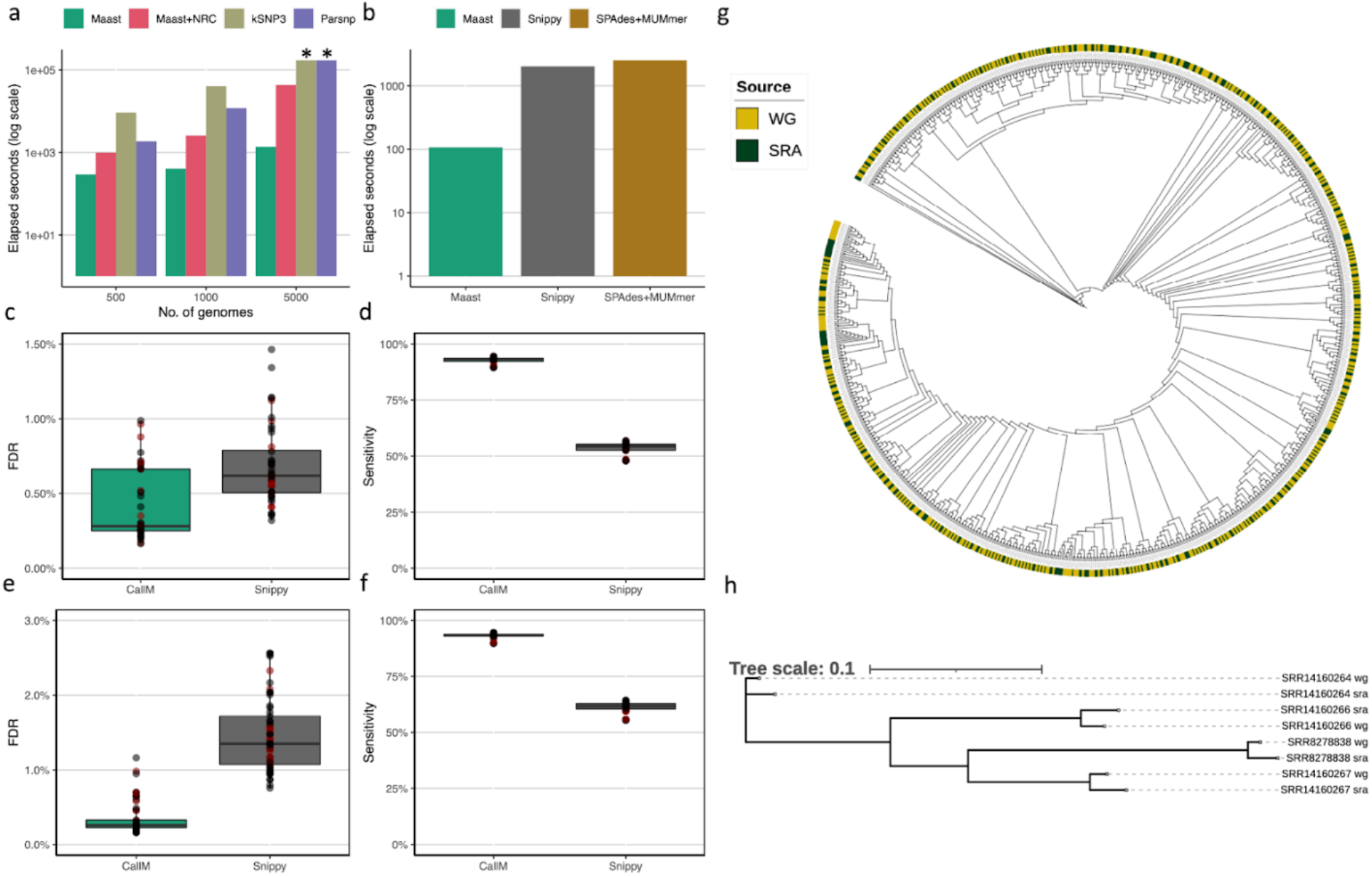
Evaluation of Maast computational performance and accuracy. (a) Comparison of genome genotyping speed between Maast, Maast without redundancy collapsing (Maast+NRC), kSNP3 and Parsnp. All methods were run on 500, 1000 and 5000 simulated *A. rectalis* genomes. (b) Comparison of short-reads genotyping speed between Maast, Snippy and SPAdes. All three methods were run on 63 strains of *B. uniformis*, whose whole genome sequencing reads (~150 million) were downloaded from the Culturable Genome Reference (CGR) study. The y-axis of both (a) and (b) indicates elapsed seconds of running in log scale. Fewer elapsed seconds indicates better performance (faster processing speed). (c-f) Comparison of Maast and Snippy genotyping accuracy at non-reference alleles of SNPs in the Maast SNP panel, based on short reads (c and d) simulated from isolate genomes with sequencing error (15× coverage) and (e and f) downloaded from isolate whole-genome sequencing projects. Both Maast and Snippy were run with default settings. (c and e) False discovery rate (FDR) comparison, where false discoveries are genotype calls that do not match the genome. (d and e) Sensitivity from the simulations in (c) and downloaded reads in (e). Sensitivity is the probability of detecting genotypes present in the genome. Color of points in c-f indicates whether the data comes from tag genomes (black) or not (red). Samples colored in red is regarded as novel to Maast databases. (g and h) Maast genotype concordance between (g) genome and short reads or (h) genome and long reads. In (g), strain population structure of *H. pylori* was reconstructed using SNPs from 473 strains. Each strain has a whole genome sequence (WG) and a short read sample (SRA) as indicated in the stacked color rings. In (h), the population structure of *H. pylori* was reconstructed from 4 strains with whole genome sequence (WG) and long reads (SRA). * kSNP3 and Parsnp runs > 48 hours on 5,000 genomes and were manually terminated, and we plot a runtime of 48 hours with the note that no output was produced.

We also evaluated the computational performance of Maast when calling SNPs in WGS reads. As a benchmark, we selected 63 whole genome sequencing samples of *Bacteroides uniformis* from the Culturable Genome Reference (CGR) study (Methods, Table S4, mean=2.38 million reads per sample). Performance was compared with Snippy as well as an assembly based strategy using a combination of SPAdes and MUMmer. Maast was ~19-24 fold faster than these methods (Fig. 3b, d) taking <2 seconds to process each WGS sample and requiring 11.8 GB RAM. Altogether, we conclude that Maast greatly accelerates SNP genotyping in both assemblies and short reads and is friendly to personal computing devices.

### Maast SNP genotypes are highly accurate

To validate the accuracy of Maast SNP genotypes, we first compared Maast to Parsnp and kSNP on 1,000 *A. rectalis* genomes with simulated SNPs (Methods). Erroneous alignments can in theory produce both incorrect genotypes (false positives) and missing genotypes (false negatives). False positives were only observed for kSNP (Table S5) while false negatives were very low for Maast (n=1) and Parsnp (n=4) but much higher for kSNP (n=5,073). These results suggest that Maast is at least as accurate as currently available methods.

To evaluate the genotyping accuracy of Maast on short reads, we compared Maast to Snippy by running both tools with their default settings on Illumina short reads simulated from 45 isolate genomes of a single species (*B. uniformis*, Table S4, 15× coverage per genome). For each method, predicted genotypes were compared to a set of ground truth genotypes determined from whole-genome alignment of the 45 isolate genomes to the Maast reference genome. Across the 45 isolates, the median SNP false discovery rate (FDR) was very low for both methods (0.28% and 0.62% for Maast and Snippy; Figure 3c and S7a). Meanwhile, the sensitivity of Maast was consistently higher (median=93.2%) compared to Snippy (median=54.3%) (Figure 3d and Figure S7c), since Maast is less subject to false negatives due to reference bias[13] and coverage filtering (i.e., minimum 1X by default compared to minimum 10X for Snippy). Since simulated reads may capture less contamination and sequencing errors than real WGS data, we repeated the evaluation with 63 short read libraries of the same species (*B. uniformis*, Table S4) downloaded from the CGR study. Similarly, we observed that Maast is more accurate in terms of both FDR (median=0.26%) and sensitivity (median=93.3%) compared to Snippy (1.35% and 61.7%) (Figure 3e, f and S7b, d). We also noted that the FDR for Maast was similar to that on simulated reads, whereas it doubled for Snippy. We conclude that Maast is a more accurate and sensitive tool compared to Snippy, reflecting advantages of 31-mer exact matching over read alignment for SNP genotyping, especially for sequencing data with relatively more contamination or sequencing errors.

### Maast reveals global genetic structure of 3,178 H. pylori isolates

To demonstrate its scalability and utility, we leveraged Maast to analyze 3,178 *H. pylori* wholegenome sequences from 39 countries across six continents (Fig. 4a, S8, Table S7). These included isolates from five animal species in addition to humans (Fig. S9). Overall, we identified 74,962 common SNPs in the core genome of *H. pylori* using Maast (Methods), which is >10 times more than a recent study [16]. Using these genotypes, we measured genetic distance between strains across continents and by host species, sex, and disease status (Fig. 4b, S10, S11). We observed clear associations between *H. pylori* genotypes and geography (Fig. 4b and c) which supports the finding that *H. pylori* has distinct populations across the globe [17]. Within the same continent, *H. pylori* strains from the same host species are more similar than those from the different host species (p-value < 2.2e-16, Wilcoxon rank sum test). Interestingly, we observed *H. pylori* strains from different host species but the same continent are more similar than those from human hosts living in different continents (Fig. 4e). This is likely due to *H. pylori* populations being highly heterogenized across continents, suggesting geography is a transcending factor over host species. We also observed that genetic distances between *H. pylori* strains are on average lower (p-value = 2.27e-13) between pairs of healthy human hosts compared to diseased pairs (Fig. 4f and S12), implying greater genomic commonality between non-pathogenic *H. pylori* strains.

**Figure 4.**
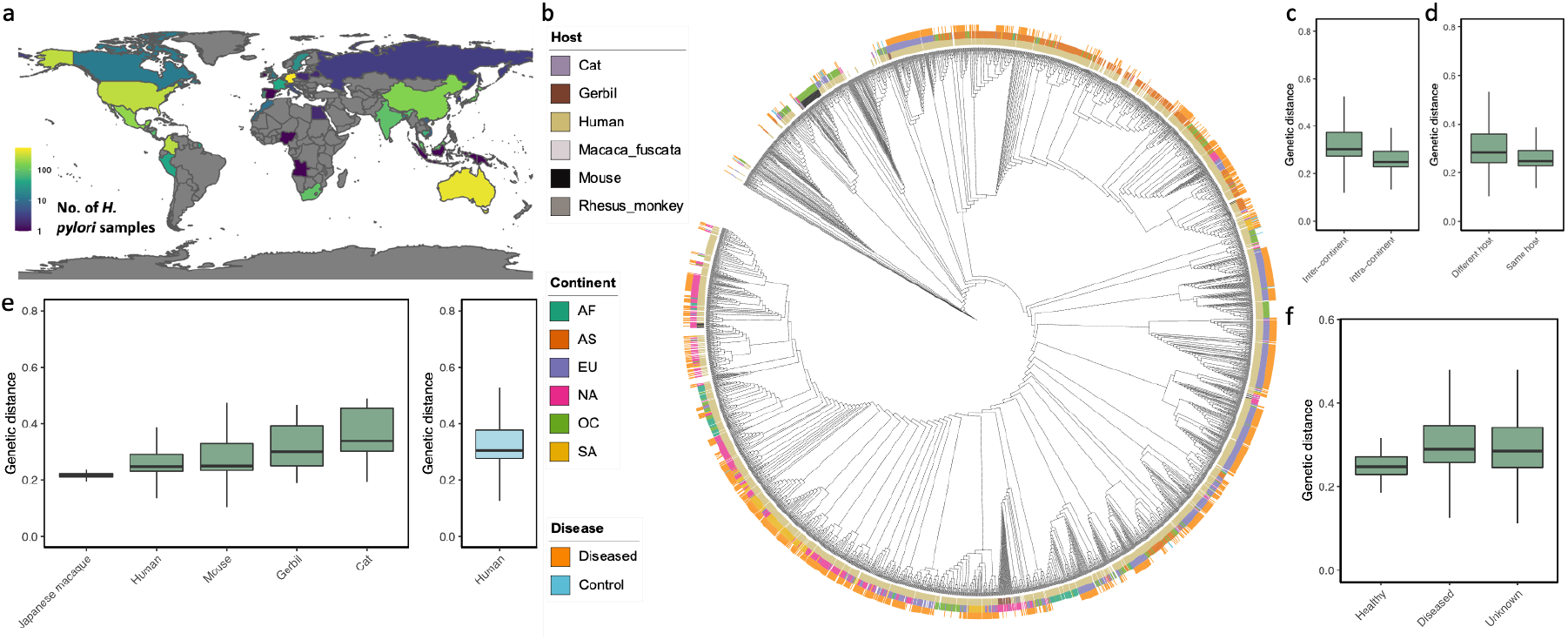
Global genetic structure of *H. pylori* genomes. (a) Geographic distribution of 3,068 *H. pylori* strains across 39 countries with color indicating the number of strains from each country. (b) Strain population structure of *H. pylori* strains reconstructed from their Maast SNP genotypes. Stacked color rings indicate the host species (inner), continent (middle), and human host disease status (outer). (c-f) Comparison of genetic distances between pairs of strains from (c) same versus different continent, (d) same versus different host species within the same continents, (e) human versus other host species and (f) control vs diseased human hosts. In (e), genetic distances are calculated between pairs of strains from the same continent where one is from a human host and the other is from the indicated host species (left) or between pairs of strains from human hosts from different continents (blue box at right). The *H. pylori* strains that infected Rhesus monkey were inoculated [32] and thus excluded.

Next, we analyzed 478 *H. pylori* isolates downloaded from NCBI (Table S8) to assess whether we would obtain the same results using Maast with unassembled sequencing reads instead of whole genome sequences (Methods). For these strains, we observed the expected similarity between isolates and their unassembled WGS reads, except for a few highly similar isolates which cannot be distinguished based on common SNPs (Figure 3g and S13). Next, we extended this analysis to four distinct *H. pylori* strains that have both whole genome assembly and long read data. We found perfect clustering of assemblies with the long-read data from the same strain (Figure 3h). Altogether our evaluation suggests that the two Maast genotyping workflows, although using distinct algorithms, are highly concordant. This means that genomic data from different sources can be pooled for analysis with Maast without assembly or other preprocessing.

### Maast enables fast tracking of SARS-CoV-2 variants

To apply Maast for global pathogen surveillance, we analyzed 37,096 *SARS-CoV-2* (SC2) strains, including 8,734 isolate genome assemblies and 28,362 WGS read samples, collected from 60 countries over a span of ~200 days (downloaded from NCBI on July 19, 2020, Table S9). Maast was able to process the data in <9h with 36 threads and a peak RAM use of 20 GB on an AWS EC2 instance (r5.16×large). Analyzing patterns in these genotypes, we first observed a clear divergence of SC2 strains relative to strain NC_045512 which was one of the first sequenced SC2 strains (Fig. 5a). Further, we found that genetic distance between SC2 strains was slightly lower within countries than between countries (Figure S16). Within-country genetic distance differed across countries, likely due to differences in temporal adjacency of samples (Fig. 5b and S17). However, we also observed exceptions where geographically adjacent countries with similar temporal sampling patterns have different levels of within-country genetic distance, such as Poland and Germany (Fig. 5b and S17), suggesting other underlying factors such as single versus multiple introduction sources. To track the differentiation of SC2 strains, we next used SC2 SNPs to perform dimension reduction, and observed changing subspecies genetic structure over time with the emergence and fading of SC2 strain clusters (Fig. 5f). These results suggest that Maast is computationally efficient and accurate enough to be used to monitor the dynamics of genetic structure of emerging pathogens at a global scale.

**Figure 5.**
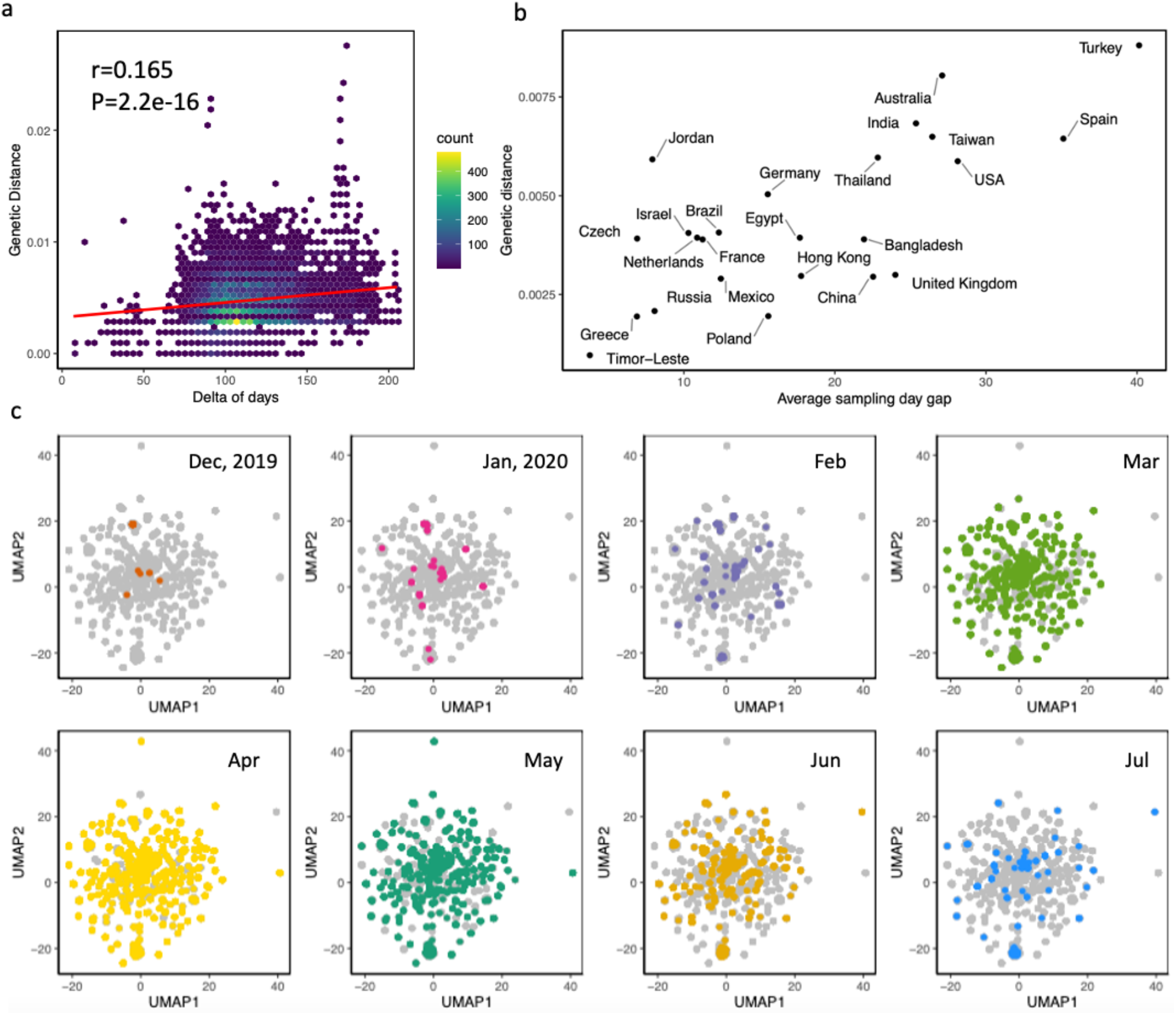
Application of Maast to track *SARS-CoV-2* diversification from Dec 2019 to July 2020. (a) Genetic distances of SARS-CoV-2 strains over time compared to one of the earliest *SARS-CoV-2* strains (Accession #: NC_045512). (b) Median genetic distance of *SARS-CoV-2* in a country is strongly correlated with average sampling day gap. (c) Subspecies structure of *SARS-CoV-2* over time. Each dot is a *SARS-CoV-2* strain, colored by sampling month with other time points in grey. Dimension reduction and visualization performed with UMAP. Nearby samples in UMAP space have similar genotypes.

## Discussion

In this study, we presented Maast, a new software tool for discovering SNPs in conspecific genomes that also includes pipelines for genotyping the panel of discovered SNPs in sequencing reads (long or short) and genome assemblies. In terms of SNP discovery, Maast is faster than existing methods due to our DynCC algorithm that identifies a subset of diverse genomes that can be used for the computationally intensive step of comparing all pairs of genomes without losing sensitivity. In fact, using these tag genomes improves our ability to identify SNPs at a given MAF and prevalence in the broader population by reducing bias due to uneven sampling of sequenced genomes. In terms of SNP genotyping, Maast uses less RAM and disk than existing methods while being about two orders of magnitude faster due to a combination of efficient data structures and novel genotyping algorithms for both genomes and unassembled reads.

The computational efficiency and accuracy of Maast enables a broad array of potential applications. In this study, we demonstrated how Maast can be used to study the genetic structure of *H. pylori*, a bacterial species with strong biogeographic patterns. We also used the early months of the *SARS-CoV-2* pandemic as a case study to illustrate that Maast can be applied to emerging viral pathogens. Maast’s ability to process thousands of genomes make it a useful tool for tracking microbial evolution in real time. By constructing and maintaining a SNP panel and k-mer database for any medically important species over time, Maast can be applied to rapidly genotype thousands of new samples without the need for genome assembly, enabling new variants to be monitored as they spread. Beyond these applications, Maast is also ideally suited for population genetic investigations of any species with many assembled or unassembled genome sequences. Its genotypes can be utilized to measure rates and patterns of selection, recombination, drift, and migration amongst lineages of a species.

Despite its advantages, Maast has several limitations. One challenge for SNP discovery is that Maast, like Parsnp and other methods requiring whole genome sequences, cannot be applied to species with very few high-quality assembled genomes. This is a common scenario for uncultured prokaryotic species plus many eukaryotes and viruses beyond the best studied pathogens, although metagenome-assembled genomes and single-cell sequencing methods are closing this gap [18–20]. The main trade-off of Maast’s genotyping algorithm is that it uses SNPs discovered in an initial collection of assembled genomes and hence may fail to detect novel variants in short reads or genomes, including rare variants and SNPs that are common in lineages not well-represented in the SNP panel. Nonetheless, as high-quality near-complete genome sequences become available for more diverse members of many species, we expect Maast will enable further applications to uncover the vast genetic structure of earth’s microbiomes.

## Conclusions

Maast is an open-source software tool that provides a highly efficient method for discovering and genotyping SNPs using either whole-genome sequences or unassembled sequencing reads. It includes a novel algorithm that dynamically collapses thousands of genomes into a diverse, representative set of tag genomes that can be used to discover most common SNPs with reduced computational resources and less bias than using all genomes for a species. This SNP discovery method is combined with a genotyping strategy that is a hybrid of whole-genome alignment and k-mer exact matching. Maast’s genotyping method achieves higher speed and accuracy than existing tools, scaling up population genetic analysis to an unrivaled number of strains compared to state-of-the-art methods. We have demonstrated that Maast can efficiently reconstruct the genetic structure of *H. pylori* and track *SARS-CoV-2* variants during the COVID-19 outbreak, applications that involve thousands of strains across the globe. As the number of species with extensive genome sequencing continues to grow, Maast is poised to catalog this vast genetic variation.

## Methods

### Maast Overview

Maast is an open-source bioinformatics tool for fast and accurate SNP genotyping from conspecific genome assemblies and sequencing reads. The development of Maast is motivated by the following observations: 1) redundant genomes are common in genome databases and these contribute little to SNP discovery while imposing a computational burden, 2) the level and pattern of redundancy varies across species, 3) the choice of reference genome affects SNP discovery, and 4) for sequencing reads, genotyping by assembly first or read alignment does not scale with increasing amounts of data.

Maast takes a set of intraspecific genomes as input and has the following major steps per species: 1) estimate the pairwise genomic distance between genomes and identify genome clusters, 2) collapse clusters to a single tag genome per cluster, 3) pick a centroid genome as reference genome, 4) perform multiple whole-genome alignment, construct consensus genome and call SNPs to generate a SNP panel, and 5) leverage the SNP panel to perform *in silico* genotyping for genome assemblies or sequencing reads.

### Pairwise genomic distance calculation

Maast estimates the genomic distance between a pair of genomes using Mash [21] (version 2.2). Mash distance can be used to approximate whole-genome average nucleotide identity (ANI) between similar genomes (e.g., conspecific genomes). For input genomes, Maast first builds 20-mer profiles or Mash sketches using its “sketch” subcommand with default parameter except for option “-s 5000” (Figure S1). Next, Maast calls the “dist” subcommand in Mash with default options to compute pairwise genomic distances.

### Identify genome clusters and tag genomes

Maast automatically determines the number of genome clusters (n) based on the user-defined minor allele frequency (MAF): n = 1 / MAF. By default, SNPs are defined as genomic sites with MAF > 1%. In this case, at least 100 genomes are required to enable an effective resolution of 1%, that is 1 in 100 genomes has the minor allele. With higher MAF thresholds, fewer genomes are needed, and vice versa.

We implemented a dynamic graph algorithm (DynaCC – Dynamic Connected Component search) in Maast to identify tag genomes based on pairwise Mash distances. DynaCC starts with a complete graph where nodes are genomes and edges are weighted by distance. The algorithm applies a distance cutoff (d-cut) strategy to prune edges, where pairs of genomes (edges) with a distance higher than a d-cut are deemed sufficiently dissimilar and removed. By applying a stringent d-cut, many edges are deleted, which reduces the complete genome graph to connected components and isolated nodes. A connected component here is analogous to a genome cluster which is defined as a maximum set of nodes in which a path exists between every pair of nodes, and a path is a sequence of edges which joins a pair of nodes. The algorithm identifies tag genomes as a union of hub genomes from connected components and all isolated nodes, where a hub genome is the genome with most edges in a connected component.

Next, we sought to implement an algorithm that solves the problem of identifying ~n tag genomes from N genomes, where N > n. Since the level and pattern of redundancy within species varies, an identical d-cut can result in detected n (n’) drastically different across species. When n’ is lower than the target n, SNPs are called with a de facto higher MAF, resulting in lower sensitivity for detecting SNPs. When n’ is much higher, the benefit of computing efficiency will diminish. It is also unrealistic to predict a static d-cut per species, due to the unknown diversity of input genomes. Here DynaCC searches and determines an optimal d-cut per species dynamically with respect to the input genomes. It first uses a range factor (rf; default 1.2) designating a critical range of n (n to n*rf) that is acceptable, since the exact n may never exist in the search space. It then applies an initial d-cut (default 0.01) and if n’ is lower than n, it searches the lower bound of d-cut with an exponential decay by a factor of 10 until an n’ higher than n is found or a hard bound (default 0.0001) is reached. Between the initial d-cut and lower bound, the algorithm performs a binary search until an n’ falling in the critical range is found or the maximum searching step (default 10) runs out. When no optimal n’ is found, the algorithm settles with a suboptimal n’ which is both higher than and nearest to n. The resulting n’ tag genomes are used for the downstream workflow.

### Identify the centroid of tag genomes

The choice of reference genome is important to SNP discovery. Maast objectively picks a reference genome for each species as the centroid of tag genomes. Maast calculates all pairwise genomic distances between tag genomes using Mash. The tag genome with the lowest average distance to all other tag genomes is selected as the centroid genome for the species. By default, Maast uses the centroid genome as the reference genome for multiple sequence alignments as well as SNP calling. Unless otherwise mentioned, we used the centroid genome for each SNP analysis in this study.

### Multiple sequence alignment and SNP calling

For each species, we performed whole-genome alignment by aligning each intraspecific genome to the centroid or otherwise specified reference genome using MUMmer [22] (version 4.0.0beta2) with the default parameters. Unreliable and repeat-induced alignments were removed using the delta-filter program from MUMmer with options ‘-q -r’ and the remaining alignments were then extracted using the show-coords program with default parameters. To promote the quality of multiple whole-genome alignment, we removed regions that are overly short (< 500bp) or poorly aligned (alignment ANI < 95% of whole-genome ANI). Maast concatenates all qualified alignments and calls SNPs as sites with the following characteristics: 1) two or more nucleotides, 2) present in at least a user-supplied percentage of genomes (prevalence threshold, default >= 90%), and 3) minor allele frequency greater than a user-supplied value (default >= 1%). Finally, Maast organizes the called SNPs into a panel and output it in a standard VCF format.

### *Maast in silico* genotyping of SNPs

For genome assemblies, Maast leverages the SNP panel from upstream steps to perform genotyping. Maast aligns each query genome, which could be non-tag genomes from SNP discovery or other additional input genomes, to the centroid genome using MUMmer (version 4.0.0beta2) with consistent quality control steps and parameters identical to those used in SNP calling. For each SNP in the panel, Maast then searches all best alignments to locate the genomic position of the SNP. If the genomic position of a SNP is found in an alignment, Maast reads and reports the allele on the query genome, otherwise Maast reports the SNP as missing.

For sequencing reads, Maast uses a k-mer exact matching algorithm for efficient and accurate genotyping. Maast leverages the SNP panel to extract short unique genomic regions (k-mers) as probes for detecting alleles that distinguish highly similar genomes and uses these k-mers to rapidly genotype reads. For k-mer extraction, Maast identifies any 31-base SNP-covering k-mers (sck-mers) that cover (in any of the 31 bases) each of the SNP alleles. We chose 31 for k based on analyses in our previous study. For each SNP, Maast first extracts all possible 31-mers containing the SNP site from the representative genome (sck-mers for reference allele). Next, Maast extracts sck-mers with the alternative allele by sliding a 31-bp window across the SNP site in the multiple sequence alignment, selecting the most frequent 31-mer at each position. We then retrieved the reverse complements of all sck-mers. In this way, for every SNP site there will be up to 62 sck-mers targeting the reference or alternative allele. We use 64-bit integers to represent the sck-mers with 00 for ‘A’, 01 for ‘C’, 10 for ‘G’ and 11 for ‘T’ and discarded the sck-mers with wildcards (e.g. ‘N’). We sort sck-mers in colex order and build an *l*-index to quickly locate all sck-mers that share a given suffix by simply pointing to the first and last entries for each suffix. This is possible because sck-mers that end with a suffix *s* of length *l* will occupy consecutive entries in the colex sorted list. We empirically selected *l*=36 as default. For each input read, Maast first breaks it down into 31-mers and encodes them into 64-bit integers. Our goal is to find exact matches of these 31-mers with sck-mers and the following exact-match algorithm is implemented: 1) look up query suffix in the *l-mer* index, if found, 2) examine all sck-mer entries identified by the *l-mer* index one by one and report exact matches. After generating all k-mers in each metagenomic sequencing read, Maast recruits an L-bit index (L-index; last L bits / suffix of encoded k-mer) to locate a bucket of pre-sorted sck-mers in the database containing all possible exact matches to the full k-mer. The algorithm invokes a sequential search for exact matches between the full k-mer and only the sck-mers in this bucket.

The output format of *in silico* genotyping is similar for both genome assemblies and sequencing reads. It uses a concise table-shaped format for its output, in which every row represents a bi-allelic SNP site. Each row has exactly 8 fields: species, SNP ID, contig, contig position, allele 1, allele 2 and coverage of allele 1 and coverage of allele 2.

### UHGG whole genome sequences and species

We downloaded genome sequences from the Unified Human Gastrointestinal Genomes [2] (UHGG) at http://ftp.ebi.ac.uk/pub/databases/metagenomics/mgnify_genomes as of September 2019. The UHGG database is by-far the largest collection of gut microbial whole genome sequences, which are originally from both isolate assemblies and metagenome assembled genomes (MAGs). The inclusion of MAGs from diverse human populations and geographic locations is critical for capturing natural genetic variation within human gut species. From the UHGG database, we selected 146 species, each with 200 or more high-quality (completeness >= 90% and contamination rate^2^ <= 5%) whole genome sequences, which account for a total of 109,365 genomes. Twenty-nine of them have more than 1000 genomes, and *Escherichia coli_D* (species id: 102506) has the most genomes (n=6,645).

### Comparing Maast to related methods

We compared Maast to several other methods in a series of simulations and data analyses designed to evaluate different aspects of computational performance and accuracy. For SNP calling in whole genome sequences, we included two methods, Parsnp and kSNP, which represents two distinct methods. Parsnp reports SNPs in the coordinates of the specified reference genome, and kSNP does not. For SNP calling in short reads, we included Snippy, which is a representative of a widely used three-step workflow, in which short reads first are aligned to reference genomes (read mapping), mapped reads are then piled up for counting coverage per site (pile-up), and SNPs are called from site coverage profiles (SNP calling). Specifically, Snippy (https://github.com/tseemann/snippy; June 2020) uses BWA mem [23] (version 0.7.17-r1188) for read mapping, samtools [24] (version 1.12) to sort and filter BAM files, and freebayes [25] (version v1.3.5) for pile-up and SNP calling. Here we chose Snippy for comparisons and analyses as it was recently reported to be the best method overall amongst methods that follow a similar strategy [26].

In all comparisons, we ran Maast, Parsnp, kSNP and Snippy with default parameters except for a flag of “-c” for Parsnp to indicate all genomes are from the same species. For all paired-end samples, we processed only forward reads (fastq 1) for the simplicity of comparison and analysis. We skipped a sample if it does not have a forward read sample as extracted from a SRA file using fastq-dump in SRA toolkit. However, we note that using the reads of both directions can effectively increase the coverage, which thus should be recommended especially for species with low abundance.

### Computing performance evaluations

We compared Maast with kSNP3 and Parsnp to evaluate its computing performance on SNP calling with whole genome sequences. We downloaded a total of 5,214 high-quality genomes for one of the most well-sequenced bacterial species, *Agathobacter rectali*, from UHGG genome colletion (Table S2). We randomly sampled 500, 1,000 or 5,000 genome sequences and ran all three methods on these genome sequences to assess the scalability of these methods. Since both kSNP3 and Parsnp failed to run through all the *A. rectalis* genomes, we downloaded high-quality genomes from UHGG for two other well-sequenced species, i.e., 3,646 genomes for *Alistipes putredinis* and 3,170 genomes for *Bacteroides_B dorei* (Table S2), and repeated the evaluation with both species to ensure the observed issue was not limited to a single species. To enable the comparison between Maast, Parsnp and kSNP as well as the performance evaluation of each method, we simulated genomes to control level of genomic divergence within a species. For the simulation, we took a random genome (UHGG id: GUT_GENOME143713) from *Agathobacter rectalis* as a simulation template and randomly selected 10,000 genomic positions on the template to be SNP sites. For each simulated genome, 10% of the SNP sites (1,000 SNPs) were randomly selected to be non-reference alleles, which we inserted *in silico*. For simplicity, we only simulated bi-allelic SNPs. To evaluate scalability of the methods, we used three levels of input size (500, 1,000 and 5,000 genomes).

To evaluate SNP genotyping in short reads, we downloaded a total 63 samples of short reads for an arbitrary species (*Bacteroides uniformis*) from the Culturable Genome Reference (CGR) study [27]. Each sample provides a distinct sequenced strain of that species. Altogether these samples account for a total of ~150 million reads.

We evaluate computing performance of Maast and other tools on an AWS EC2 instance with the following specifications: AWS r5.16xlarge, 32 physical CPU cores (64 vCPU), Intel 8175M CPU @ 2.50GHz, 512 GB RAM and EBS gp2 RAID array providing 13,600 Mbps bandwidth. We measured both speed and peak RAM consumption using GNU time (version 1.7) command with option “-v”.

To ensure a fair comparison between different methods, all methods were run on all of the input data with the same reference genome whenever one is used. Each method was run using all cores of an environment whenever possible.

We manually terminated the running of any tool after both tools had been running over a designated maximum running time window (48 hours) and had not finished. When terminated, we estimated the speed of Parsnp and kSNP with the maximum time use of 48 hours and projected peak RAM use linearly as a function of the number of genomes.

### Accuracy evaluation of SNPs from simulated and downloaded reads

To evaluate the accuracy of SNP genotyping by Maast in short reads and to compare it to Snippy, we simulated reads from whole genome sequences. For these simulations, we downloaded a total of 45 isolate genomes for a species (*Bacteroides uniformis*) from the CGR study. These genomes were cultivated from fecal samples of healthy humans and characterized as non-redundant and high-quality draft genomes. We used InSilicoSeq [28] (version 1.4.2) with the options “--model HiSeq” and “--n_reads 2000000” to simulate reads with Illumina length and error characteristics. This generated two paired-end read files each containing ~1 million 126bp-long reads from each genome. For simplicity, we proceeded only the forward reads. We used genome coverage of 15× for simulations by randomly drawing reads from the simulated metagenomes. The number of reads required for a level of coverage c was estimated by the following formula: number of reads = c * genome length / 126. For example, to provide a 15× coverage for a genome with a size of 6M bp, a rough number of 714,285 (15 * 5,000,000 / 126) 126-bp reads are needed. To complement the simulations, we extend the accuracy analyses to the 63 samples of short reads used in the computing performance evaluation.

To ensure a fair comparison, we ran Maast and Snippy on both the simulated and downloaded reads with default parameters and the same reference genome which was arbitrarily selected. Our evaluations focused on correct identification of SNPs and on the accuracy of the alleles in the typed SNPs. For each strain, ground truth genotypes are determined by aligning the whole sequence of that strain to the reference genome. We focused our evaluations on the SNPs that are potentially able to be genotyped by both Maast and Snippy, i.e., SNPs in the Maast panel. True positives (TP) and false positives (FP) are the correct and incorrect genotypes compared to ground truth; false negatives (FN) are sites with no genotype or an incorrect genotype. In this way, we conveniently calculate the false discovery rate (FDR) as 1 minus the ratio between the sum of TP sites and the sum of all reported sites and the sensitivity as TP/(TP+FN) or the ratio of TP sites to all Maast sites. These values were calculated for each method on non-reference sites only as Snippy only reports them.

### Reconstruction of genetic structure of H. pylori

To show Maast is powerful for exploring genetic structure within microbial species, we downloaded a total of 3,524 whole genome sequences of *H. pylori* from NCBI and PATRIC [29] website as of March 2021. We kept 3,428 high-quality genomes (completeness >= 90% and contamination rate <= 5%) as determined using checkM, which include 1,672 and 1,756 genomes from NCBI and PATRIC, respectively (Table S6). We ran the SNP calling module of Maast with default parameters (site prevalence > 0.9 and MAF > 0.01) on the whole genome sequences, resulting in 107 tag genomes and a panel of 275,116 SNPs. We then ran the database building module of Maast with default settings and generated a sck-mer database with 3,566,180 sck-mers and 74,962 SNPs that are covered by at least one sck-mer.

Next, we sought to determine whether the common SNPs discovered and typed by Maast could be used for biogeographic analyses of diverged *H. pylori* strains. To maximize the number of strains in this analysis, we downloaded an additional 1,522 short read samples from NCBI SRA as of April 2021, each representing a distinct strain. We then combined these short-read samples together with 1,756 PATRIC genomes to form one of the largest *H. pylori* strain collections (n=3,178) to date with diverse host and geographic information (Table S7). We did not include NCBI whole genome sequences due to less richness in metadata. We ran Maast on these strains and used the RAxML [30] algorithm to calculate pairwise genetic distance as well as construct a phylogenetic tree based on concatenated alleles of SNP sites genotyped. We uploaded the resulting RAxML tree file to the iTol [31] website for visualization. To account for possible bias due to less covered SNPs, we applied two filters to both the SNPs and strains: 1) we excluded a SNP from this analysis if it is present in fewer than 5 strains, and 2) we excluded a strain if it has <1000 genotyped SNPs or < 50% of total SNPs genotyped.

To further validate consistency between Maast SNP genotyping of whole genome sequences and short reads, we identified 478 *H. pylori* strains that have both raw whole-genome sequencing reads and assembled genomes available in NCBI records (Table S8). We used similar steps to run Maast on these strains and reconstructed a phylogenetic tree.

### Tracking early outbreak of SARS-CoV-2

We analyzed SARS-CoV-2 strains from the early outbreak. In contrast to the long evolutionary history of *H. pylori*, genomic tracking of SARS-CoV-2 is quite recent and extensive, which provides a good example to evaluate how well Maast can be used to track the genetics of a burst of highly similar genomes. We downloaded a total of 37,096 SARS-CoV-2 strains from diverse geographic regions (Fig. S14), including 8,734 whole genome sequences from the NCBI *SARS-CoV-2* Data Hub as well as 28,362 short read samples available from the NCBI SRA as of July 2020 (Table S9). As expected, the Mash distances between SARS-CoV-2 genomes are extremely low due to the short evolution history (Fig. S15). To effectively differentiate these strains, we included rare SNPs in this application by running Maast with site prevalence > 0.95 and MAF > 0.001 on the whole genome sequences of SARS-CoV-2, which resulted in 1,077 tag genomes and a panel of 1,114 SNPs. With Maast, we built a sck-mer database, which includes 128,416 sck-mers and 1,045 SNPs that are covered by at least one sck-mer. Then, we genotyped SNPs for the rest of the strains with Maast and computed SNP-based genetic distance and a RAxML tree as in the *H. pylori* analysis.

To visually track the genetic change of SARS-CoV-2 over time, we identified the major allele (allele with the highest frequency across strains) of each SNP, generated a binary matrix of major allele presence/absence per strain, and performed dimension reduction on this matrix with UMAP (umap package version 0.2.3.1) in R. We plotted all strains in the resulting UMAP coordinates. We visually identified 1,073 outliers (<3% of total strains) and removed them from the plot. To account for possible bias due to less covered SNPs, we applied two filters to both genomes and metagenomes: 1) we excluded a SNP in both genomes and metagenomes if it is present in fewer than 5 metagenomes, and 2) we excluded a metagenome if it has <1000 genotyped SNPs or < 50% of total SNPs genotyped.

## Data availability

All described datasets are publicly available through the corresponding repositories. Genome assemblies and whole genome sequencing samples are available at UHGG, NCBI and PATRIC database with accession numbers in supplementary tables: genome assemblies for *Alistipes putredinis, Bacteroides_B dorei* and *Agathobacter rectalis* (Table S2), and both genome assemblies and whole genome sequencing samples for *Bacteroides uniformis* (Table S4), *Helicobacter pylori* (Table S6-S8) and SARS-CoV-2 (Table S9).

### Software availability

Maast is written in C++ and Python, and it is released as open-source software under the MIT license. The source code and documentation of Maast is available on GitHub (https://github.com/zjshi/Maast).

## Acknowledgement

This work was funded by the Chan Zuckerberg Biohub, Gladstone Institutes, and NSF grant #1563159.

## Competing interests

The authors declare no competing interests.

## Notes

### Competing Interest Statement

The authors have declared no competing interest.

### Summary of Updates

Manuscript file was modified to include an acknowledgement section to acknowledge the funding agencies and grants that supported this work.

