## Supplementary figures for "Maast: genotyping thousands of microbial strains efficiently"

#### Supplementary Figure legends

**Figure S1. Choosing default Mash sketch size for Maast.** Violin plot shows the distribution of Spearman correlations between whole genome ANI and Mash distance across 146 bacterial species. Correlation is calculated between all pairs of genomes in each species and the median is plotted. Three different Mash sketch sizes (1,000, 5,000 and 10,000) are evaluated.

**Figure S2. K-mer exact matching to genotype sequencing reads.** (a) Scheme whereby SNP-covering k-mers (sck-mers) are extracted from multiple sequence alignments. Each colored rectangle represents a SNP site and the cross color indicates distinct alleles. (b). Conceptual workflow of the k-mer exact match algorithm in Maast. All sck-mers have been binary encoded, pooled, and sorted in Colex order. The L-bit suffixes of sck-mers is used to build an index (L-index; green) whose entries point to consecutive records in the sck-mer table (cyan). Efficient exact-matching of k-mers from metagenomes starts with extracting all k-mers present in each sequencing read (input). To determine what they are, the range of entries in the L-index corresponding to each L-bit hit is queried with an exact-match algorithm. Dashed black arrows indicate processes occurring during database development, and solid gray arrows indicate the Maast genotyping workflow.

**Figure S3. Flowchart of DynaCC algorithm for collapsing redundancy in the collection of whole genome sequences.** The workflow is presented sequentially from top to bottom in three steps (grey boxes).The algorithm (a) first determines the acceptance range of number of tag genomes using the user-defined minor allele frequency threshold and a range factor (>1), (b) determines the search space of distance cuts (d-cuts) that can produce an acceptable set of tag genomes, and (c) identifies the best d-cut that produces the number of tag genomes in the acceptance range or as close as possible using binary search (white box). The algorithm terminates without output if no d-cut can be found to produce the number of tag genomes higher than the lower bound of the acceptance range. Although unlikely, it is possible that a good d-cut is accepted in step (b), which results in a successful early exit.

**Figure S4. Intraspecific diversity varies across 146 human gut species.** For each species, intraspecific diversity is indicated by the median Mash distance between pairs of conspecific genomes.

**Figure S5. Species vary in their genomic redundancy.** (a) When the same d-cut (0.01) is applied to all 146 human gut species, the species vary substantially in their number of clusters with a single-linkage clustering strategy due to different levels of genomic redundancy. Circle size indicates the number of clusters identified by Maast. If levels of redundancy were invariable across species, the ratio of the number of clusters versus the number of genomes of each species would be constant. The opposite is observed here as many pairs of species with similar numbers of genomes have drastically different numbers of clusters. Arrows point to two example species (*Agathobacter rectalis* and *Alistipes putredinis*) with different redundancy levels. (b) Rarefaction curves of common SNP (minor allele frequency >= 1%, prevalence >= 90% across intraspecific genomes) discovery for two example species: *A. rectalis* (upper, high redundancy, 46 clusters based on Mash) and *A. putredinis* (lower, low redundancy, 2650 clusters). SNP discovery increases with the number of genomes and then levels off for both species. SNP discovery levels off much more quickly for *A. putredinis*, underscoring how genomic redundancy limits SNP discovery at a given MAF threshold. The top 1,000 genomes of each species with highest quality (i.e., completeness and contamination) were included for the rarefaction analysis. The curve is made by counting the number of SNPs that can be discovered using down-sampled sets of fewer genomes from 10 to 1,000 genomes increasing by 10 genomes at each step. Each down-sampling was repeated 10 times and the mean value is shown.

**Figure S6. Relationship between Mash distances and SNP discovery with Maast.** Distribution of intra-specific Mash distances for eight example species, including (a) *Anaerotignum sp000436415* (100177), (b) *Sutterella wadsworthensis_B* (101361), (c) *Succinivibrio sp000431835* (100412), (d) *Akkermansia muciniphila* (102454), (e) *Akkermansia muciniphila_B* (102453), (f) *Alistipes shahii* (100003), (g) *Faecalibacterium prausnitzii_K* (species id: 101300) and (h) *Phascolarctobacterium faecium* (103439). Species in the upper and bottom panel are the ones with a high and low level of tag-only SNPs, respectively. Species in the same column have a similar number of genomes. The level is estimated as a fraction of all SNPs that are discovered with tag genomes but not with all genomes.

**Figure S7. Accuracy comparison between Maast and Snippy.** Individual simulated (a and c) and (b and d) isolate sequencing samples are shown. (a and b) False positive rate at SNP sites in simulated reads (a) and isolate sequencing samples (b). (c and d) Sensitivity across the simulated reads (c) and isolate sequencing samples (d) is calculated as the probability of detecting SNPs present in the isolate genome. Only variable sites are included in the calculations.

**Figure S8. Comparison of Maast genotypes from simulated forward and reverse reads.** Concordance is shown as the Jaccard similarity between each pair of samples.

**Figure S9. Geographic distribution of *H. pylori* strains by host species.** For each host species (cat, mouse, gerbil, rhesus monkey and macaca fuscata), the number of strains is indicated (colors) across 39 countries.

**Figure S10. *H. pylori* population structure is not associated with host gender.** (a) Strain population structure of *H. pylori* reconstructed from Maast SNP genotypes. Stacked color rings indicate host gender. (b) Pairwise genetic distances between strains from human hosts that are of same or different gender.

**Figure S11. *H. pylori* strain diversity is elevated in diseased hosts.** Strain population structure of *H. pylori* reconstructed from Maast SNP genotypes. Stacked color rings indicate the disease status, the diagnosis of inflammation, ulcer and cancer of the human host of *H. pylori* strains.

**Figure S12. *H. pylori* strain diversity as a function of host species and geography.** Pairwise genetic distances between *H. pylori* strains from (a) the same or different host species, (b) human hosts from the same or different continents, (c) specific host species and (d) diseased or healthy hosts. (c and d) only included strains from the same continent.

**Figure S13. Pairwise genetic distances between *H. pylori* strains from the same and different data sources.** Two types of data source are compared, including whole genome assembly and sequencing reads.

**Figure S14. Distribution of Mash distances of 8,734 sequenced SARS-CoV-2 strains.**

**Figure S15. Distribution of SARS-CoV-2 WGS projects across countries.** Only countries with >30 projects were shown.

**Figure S16. Distribution of SARS-CoV-2 WGS projects over time in different countries.** Comparison of genetic distances between pairs of SARS-CoV-2 strains from (a) same versus different countries and (b) same versus different months.

**Figure S17. Genetic distances of SARS-CoV-2 strains across countries.** Left panel shows the distribution of sampling day of SARS-CoV-2 strains. Right panel shows the distribution of genetic distance of strain pairs in each country.

#### Supplementary Figures

Figure S1

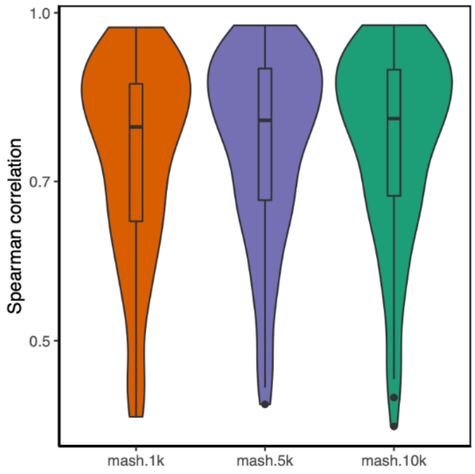

Figure S2

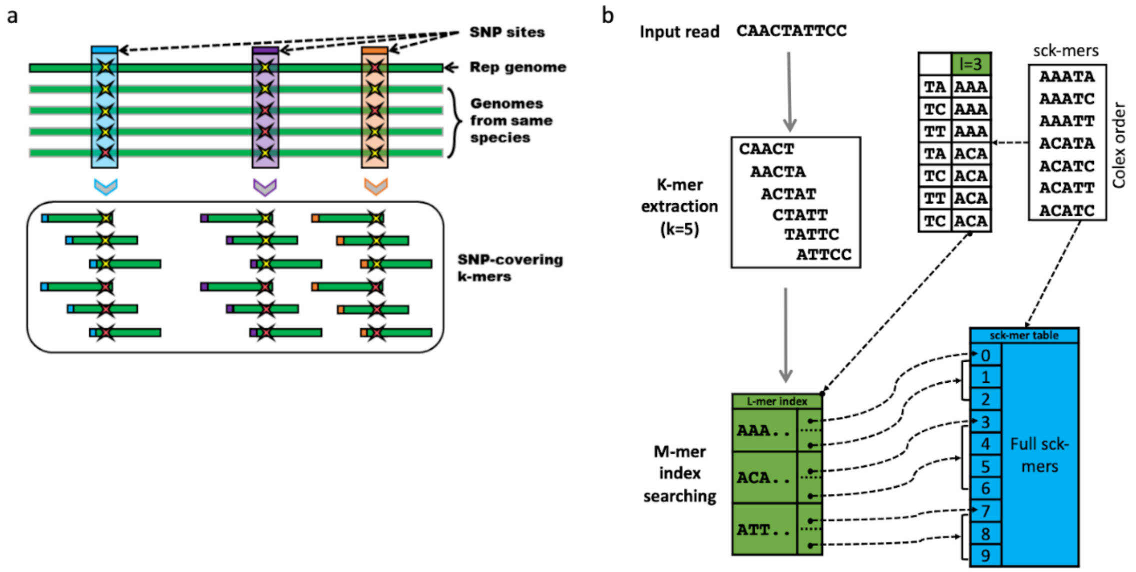
a
SNP sites
Rep genome
Genomes from same species
SNP-covering k-mers
b
Input read CAACTATTCC
K-mer extraction (k=5)
CAACT
AACTA
ACTAT
CTATT
TATTC
ATTCC
l=3
TA AAA
TC AAA
TT AAA
TA ACA
TC ACA
TT ACA
TC ACA
sck-mers
AAATA
AAATC
AAATT
ACATA
ACATC
ACATT
ACATC
Colex order
M-mer index searching
L-mer index
AAA..
ACA..
ATT..
sck-mer table
0
1
2
3
4
5
6
7
8
9
Full sck-mers

Figure S3

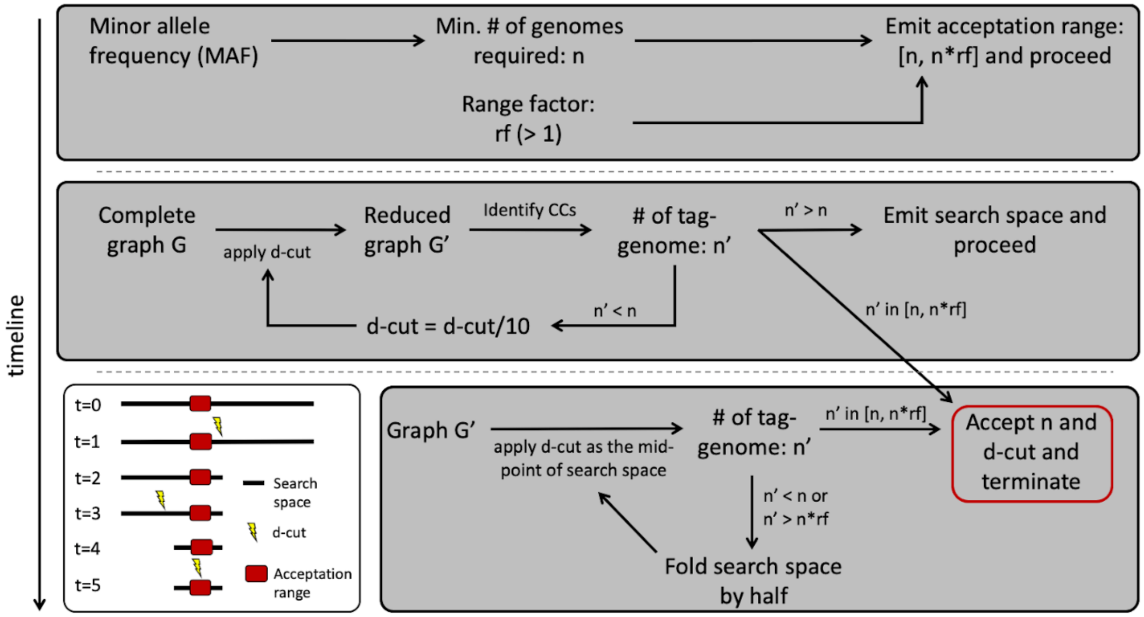

timeline
Minor allele frequency (MAF)
Min. # of genomes required: n
Emit acceptation range: [n, n*rf] and proceed
Range factor: rf (> 1)
Complete graph G
apply d-cut
Reduced graph G'
Identify CCs
### of tag-genome: n'
n' > n
Emit search space and proceed
n' < n
d-cut = d-cut/10
n' in [n, n*rf]
t=0
t=1
t=2
t=3
t=4
t=5
Search space
d-cut
Acceptation range
Graph G'
apply d-cut as the mid-point of search space
### of tag-genome: n'
n' in [n, n*rf]
Accept n and d-cut and terminate
n' < n or n' > n*rf
Fold search space by half

Figure S4

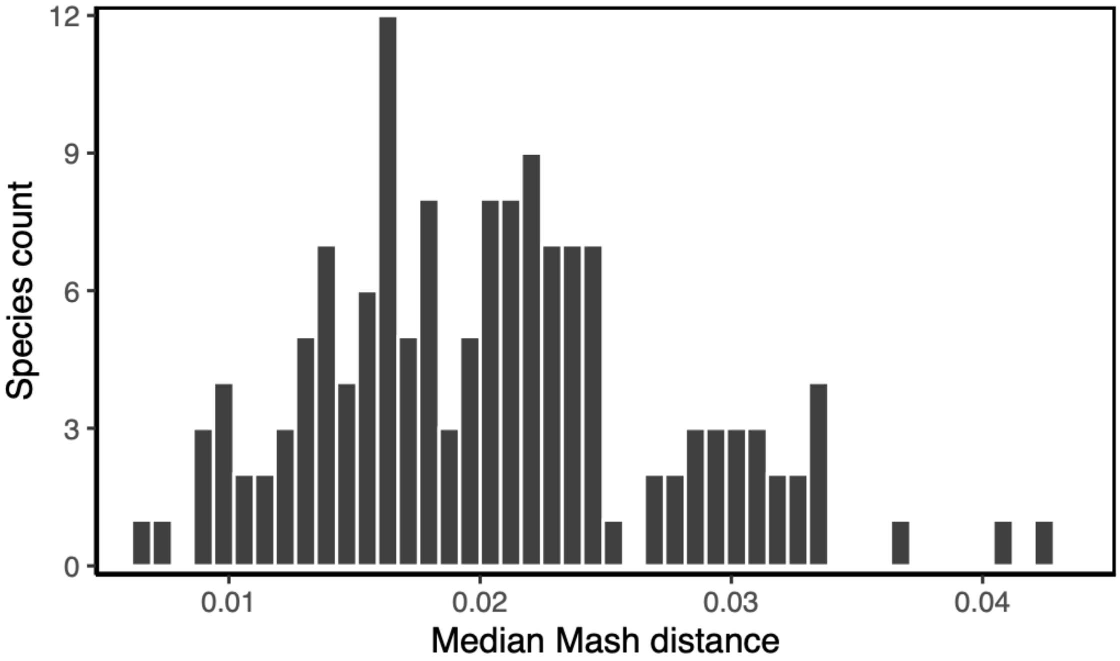

Figure S5

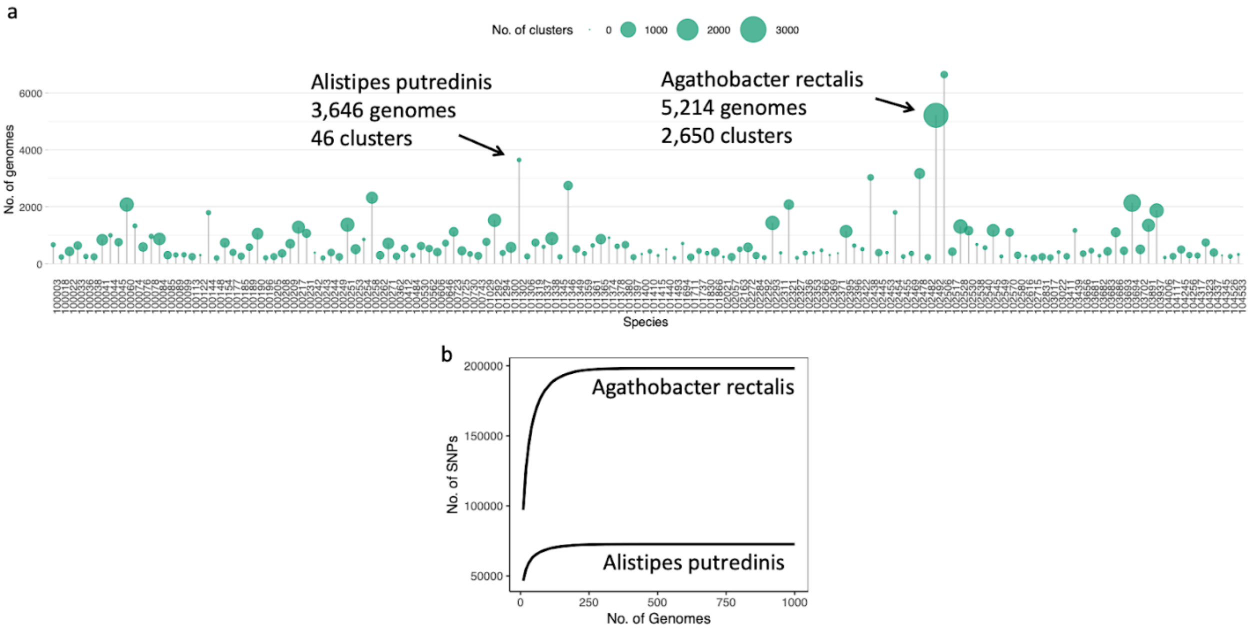
a
No. of clusters
0
1000
2000
3000
Alistipes putredinis
3,646 genomes
46 clusters
Agathobacter rectalis
5,214 genomes
2,650 clusters
No. of genomes
0
2000
4000
6000
Species
b
Agathobacter rectalis
Alistipes putredinis
No. of SNPs
50000
100000
150000
200000
0
250
500
750
1000
No. of Genomes

Figure S6

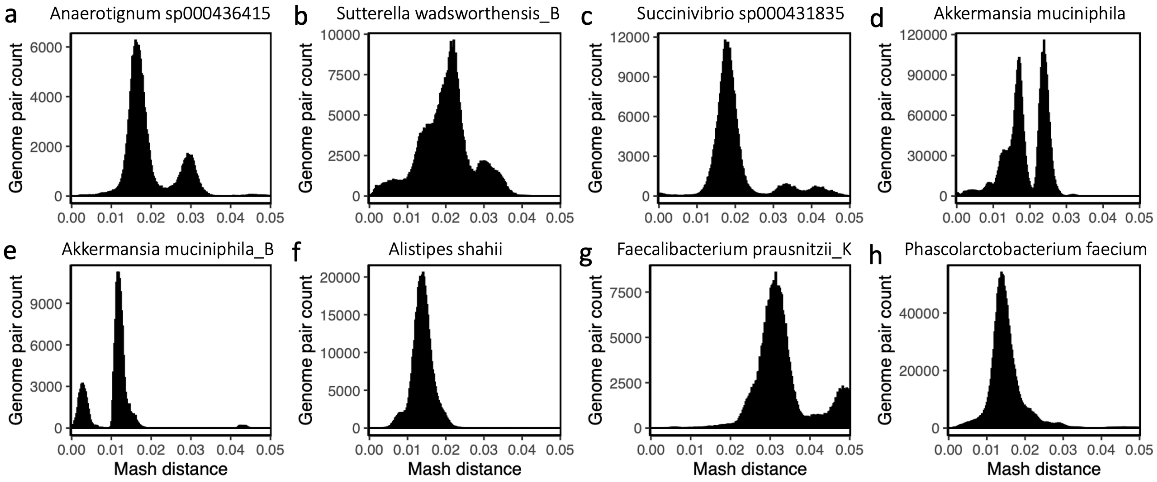

a
Anaerotignum sp000436415
b
Sutterella wadsworthensis_B
c
Succinivibrio sp000431835
d
Akkermansia muciniphila
e
Akkermansia muciniphila_B
f
Alistipes shahii
g
Faecalibacterium prausnitzii_K
h
Phascolarctobacterium faecium
Genome pair count
Mash distance
0.00 0.01 0.02 0.03 0.04 0.05

Figure S7

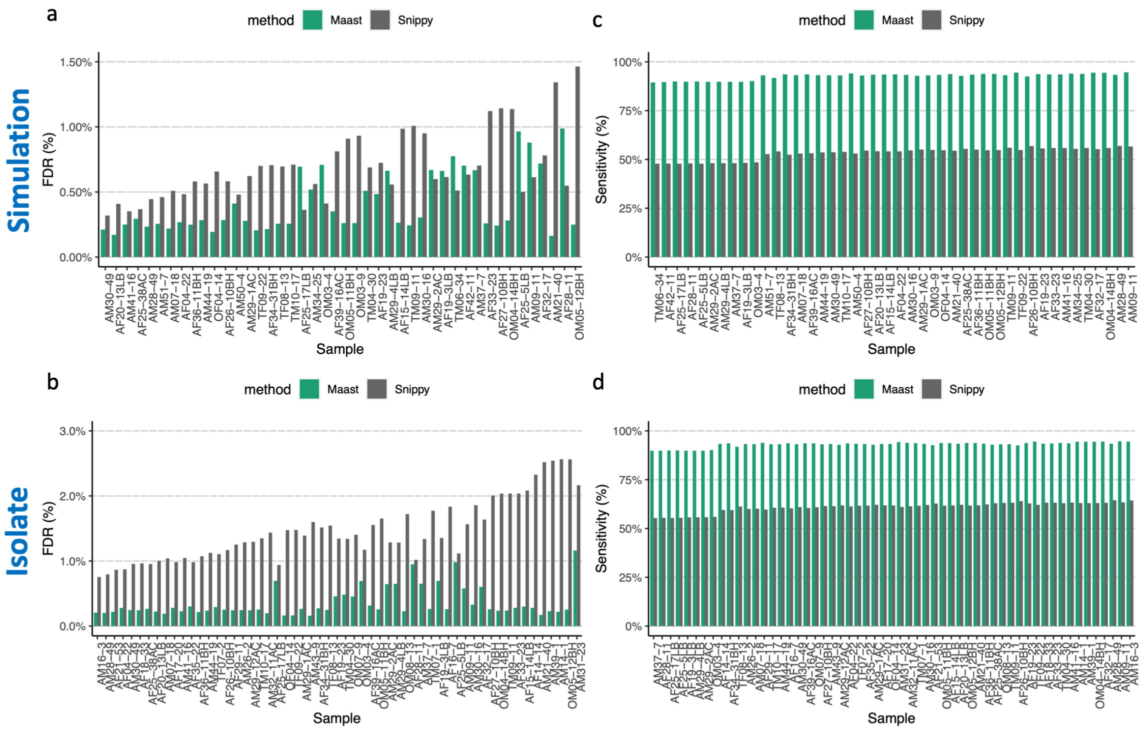

a
b
c
d
Simulation
Isolate
method
Maast
Snippy
FDR (%)
Sensitivity (%)
Sample
0.00%
0.50%
1.00%
1.50%
0.0%
1.0%
2.0%
3.0%
0%
25%
50%
75%
100%
AM30-49 AF20-13LB AM41-16 AF25-38AC AM28-49 AM51-7 AM07-18 AF04-22 AF36-11BH AM44-19 OF04-14 AF26-10BH AM50-4 AM29-1AC TF09-22 AF34-31BH TF08-13 TM10-17 AF25-17LB AM34-25 OM03-4 AF39-16AC OM05-11BH OM03-9 TM04-30 AF19-23 AM29-4LB AF15-14LB TM09-11 AM30-16 AM29-2AC AF19-3LB TM06-34 AF42-11 AM37-7 AF33-23 AF27-10BH OM04-14BH AF25-5LB AM09-11 AF32-17 AM21-40 AF28-11 OM05-12BH
TM06-34 AF42-11 AF25-17LB AF28-11 AF25-5LB AM29-2AC AM29-4LB AM37-7 AF19-3LB OM03-4 AM51-7 TF08-13 AF34-31BH AM07-18 AF39-16AC AM44-19 AM30-49 TM10-17 AM50-4 AF27-10BH AF20-13LB AF15-14LB AF04-22 AM30-16 AM29-1AC OM03-9 OF04-14 AM21-40 AF25-38AC AF36-11BH OM05-11BH OM05-12BH TM09-11 TF09-22 AF26-10BH AF19-23 AF33-23 AM41-16 AM34-25 TM04-30 AF32-17 OM04-14BH AM28-49 AM09-11

Figure S8

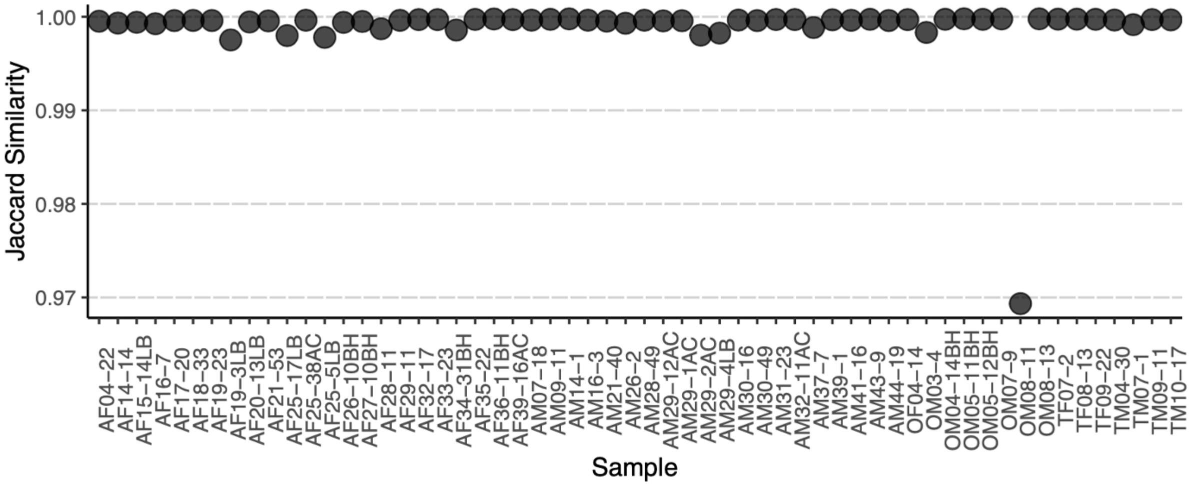

Jaccard Similarity
1.00
0.99
0.98
0.97
Sample
AF04−22
AF14−14
AF15−14LB
AF16−7
AF17−20
AF18−33
AF19−23
AF19−3LB
AF20−13LB
AF21−53
AF25−17LB
AF25−38AC
AF25−5LB
AF26−10BH
AF27−10BH
AF28−11
AF29−11
AF32−17
AF33−23
AF34−31BH
AF35−22
AF36−11BH
AF39−16AC
AM07−18
AM09−11
AM14−1
AM16−3
AM21−40
AM26−2
AM28−49
AM29−12AC
AM29−1AC
AM29−2AC
AM29−4LB
AM30−16
AM30−49
AM31−23
AM32−11AC
AM37−7
AM39−1
AM41−16
AM43−9
AM44−19
OF04−14
OM03−4
OM04−14BH
OM05−11BH
OM05−12BH
OM07−9
OM08−11
OM08−13
TF07−2
TF08−13
TF09−22
TM04−30
TM07−1
TM09−11
TM10−17

Figure S9

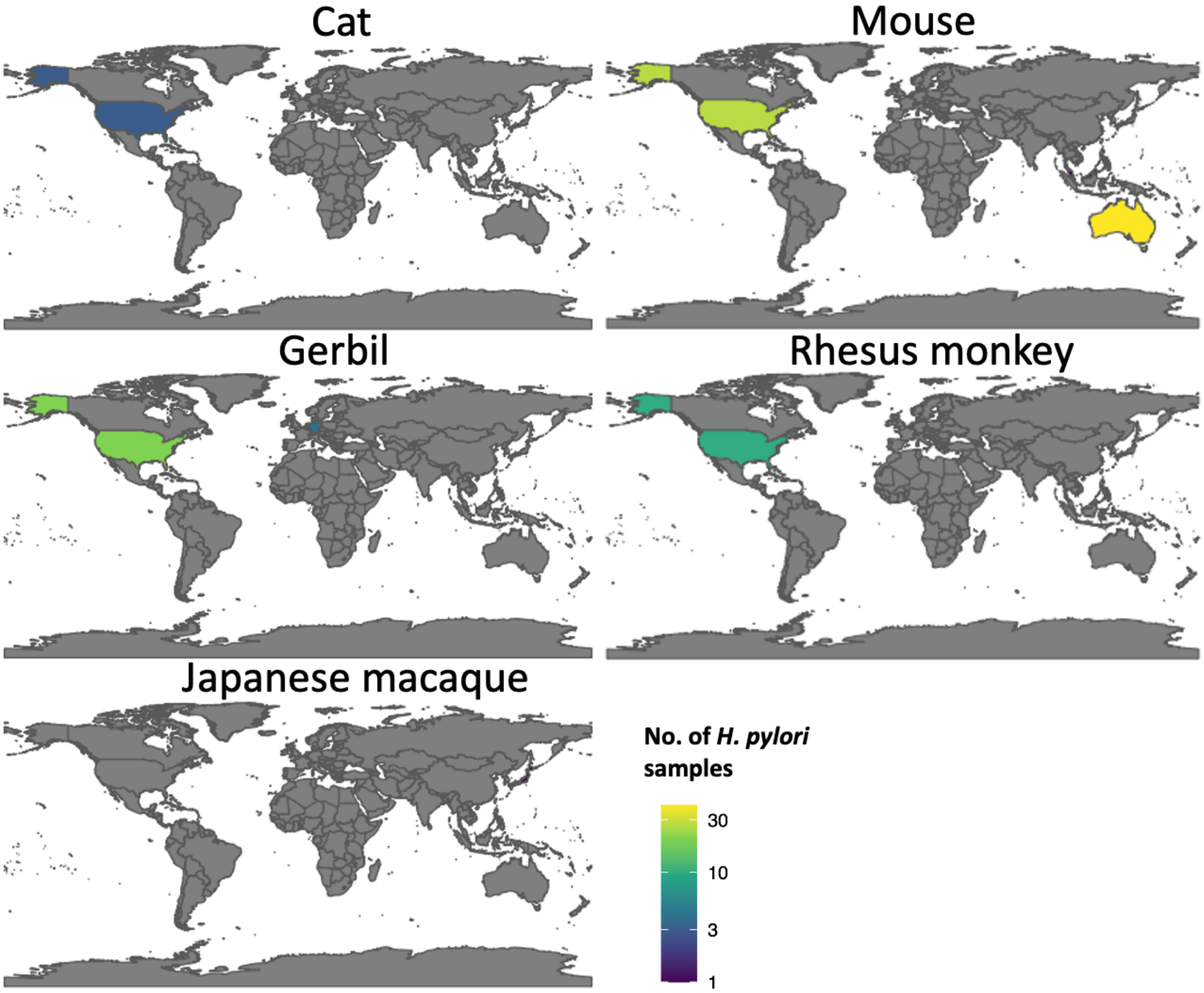

Cat
Mouse
Gerbil
Rhesus monkey
Japanese macaque
No. of *H. pylori* samples
30
10
3
1

Figure S10

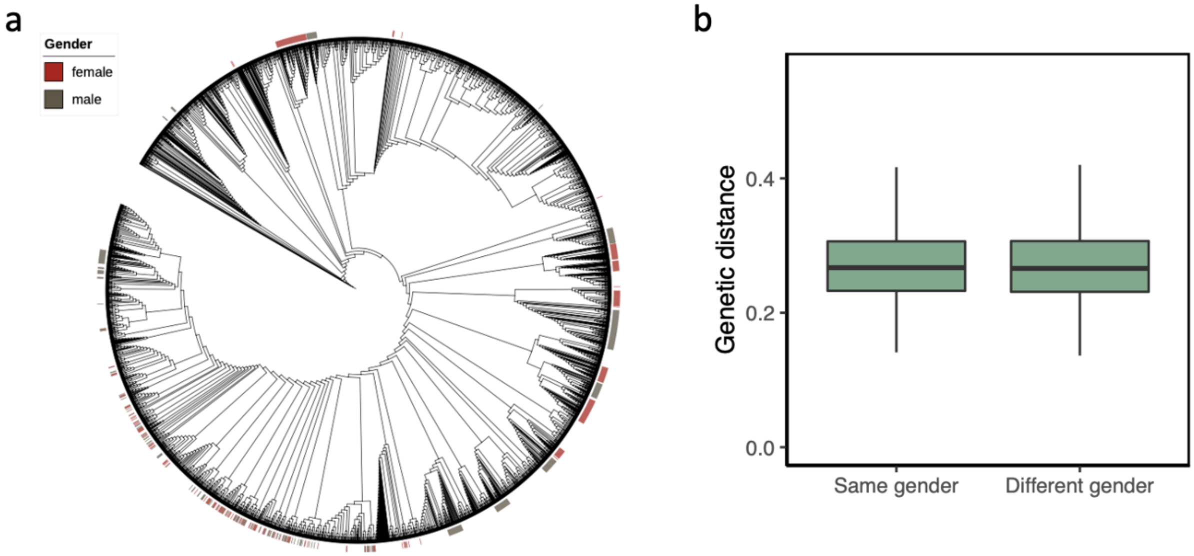
a
Gender
female
male
b
Genetic distance
0.4
0.2
0.0
Same gender
Different gender

Figure S11

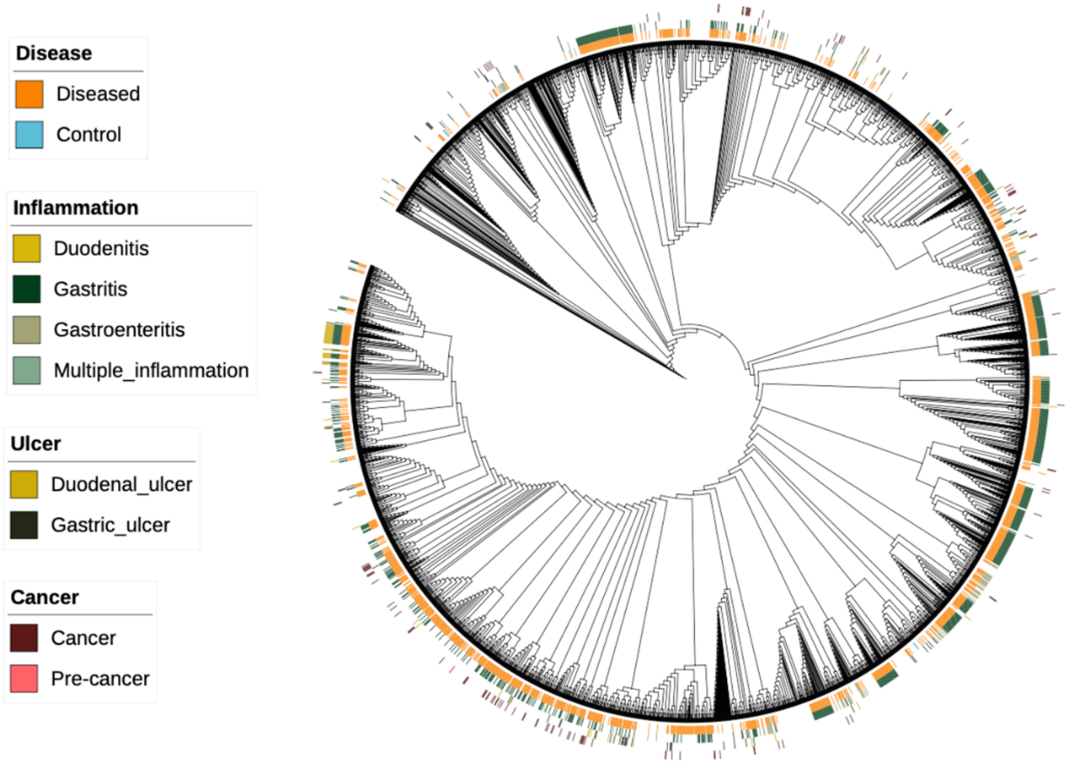

Disease
Diseased
Control
Inflammation
Duodenitis
Gastritis
Gastroenteritis
Multiple_inflammation
Ulcer
Duodenal_ulcer
Gastric_ulcer
Cancer
Cancer
Pre-cancer

Figure S12

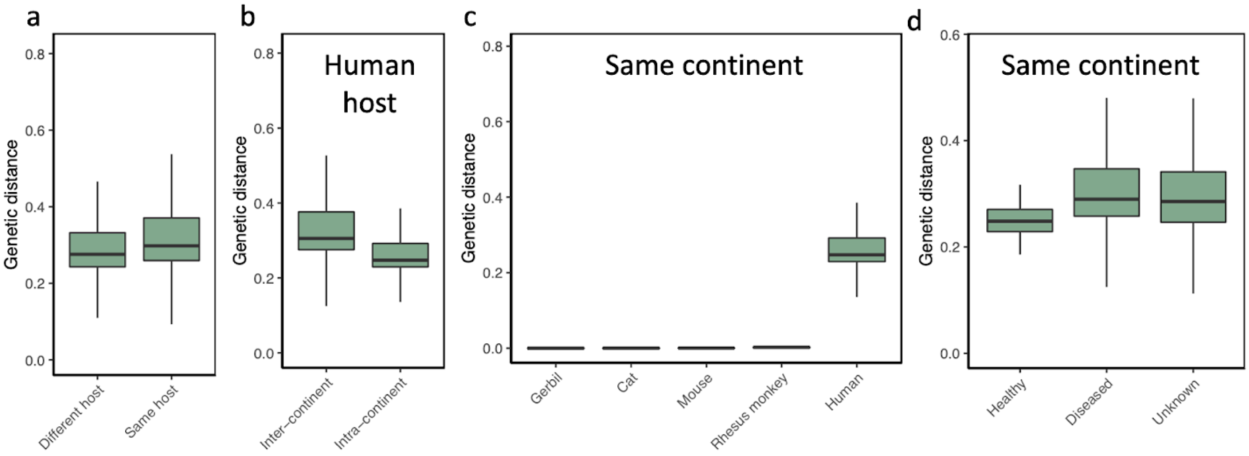

a
b
Human host
c
Same continent
d
Same continent
Genetic distance
Different host
Same host
Inter−continent
Intra−continent
Gerbil
Cat
Mouse
Rhesus monkey
Human
Healthy
Diseased
Unknown

Figure S13

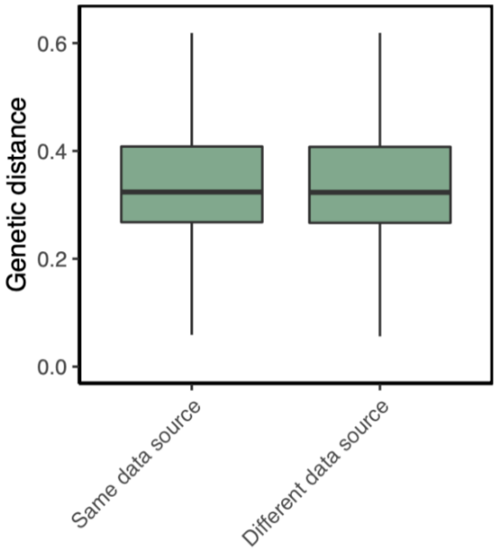

0.6
0.4
0.2
0.0
Genetic distance
Same data source
Different data source

Figure S14

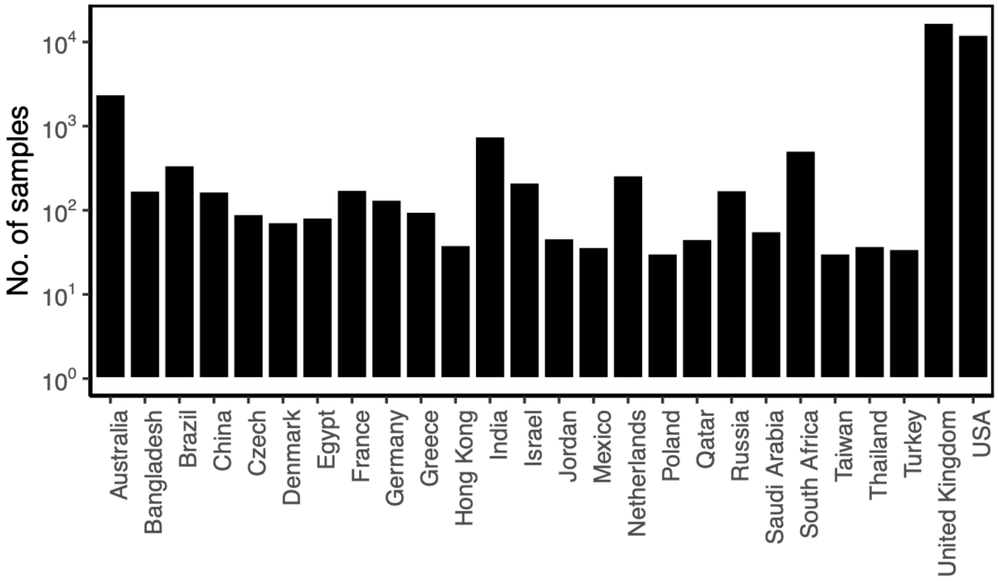
No. of samples
10^4
10^3
10^2
10^1
10^0
Australia
Bangladesh
Brazil
China
Czech
Denmark
Egypt
France
Germany
Greece
Hong Kong
India
Israel
Jordan
Mexico
Netherlands
Poland
Qatar
Russia
Saudi Arabia
South Africa
Taiwan
Thailand
Turkey
United Kingdom
USA

Figure S15

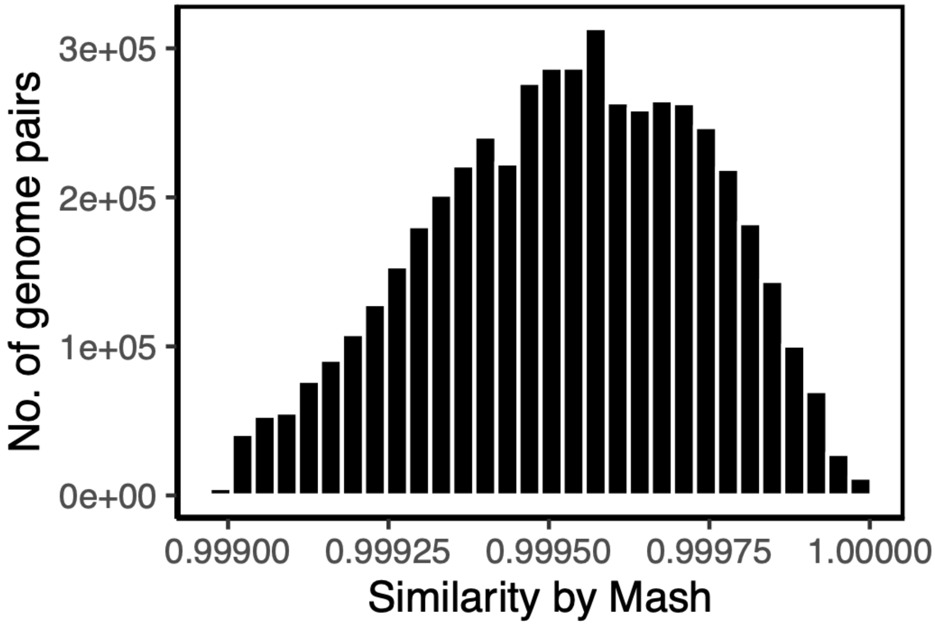

3e+05
2e+05
1e+05
0e+00
No. of genome pairs
0.99900
0.99925
0.99950
0.99975
1.00000
Similarity by Mash

Figure S16

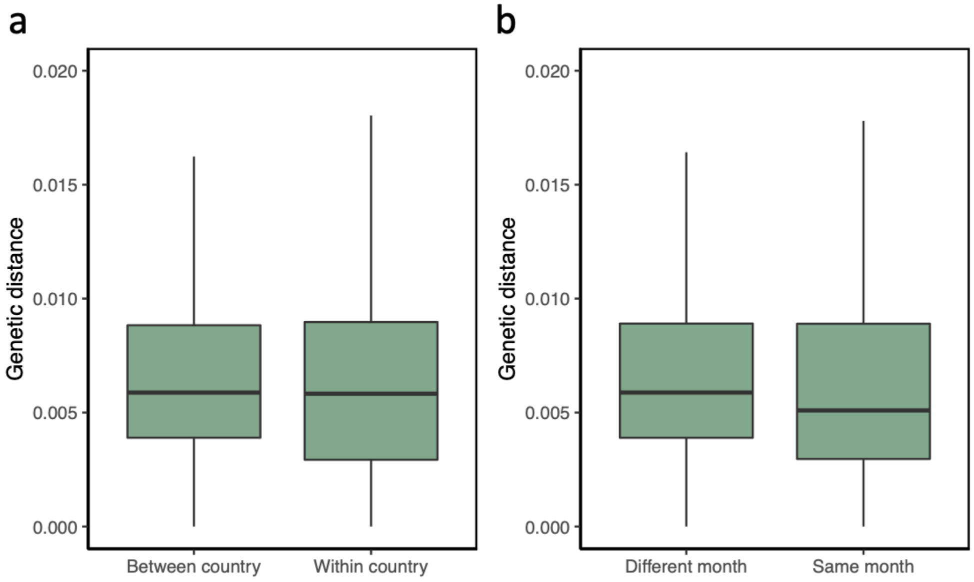
a
b
Genetic distance
0.020
0.015
0.010
0.005
0.000
Between country
Within country
Different month
Same month

Figure S17

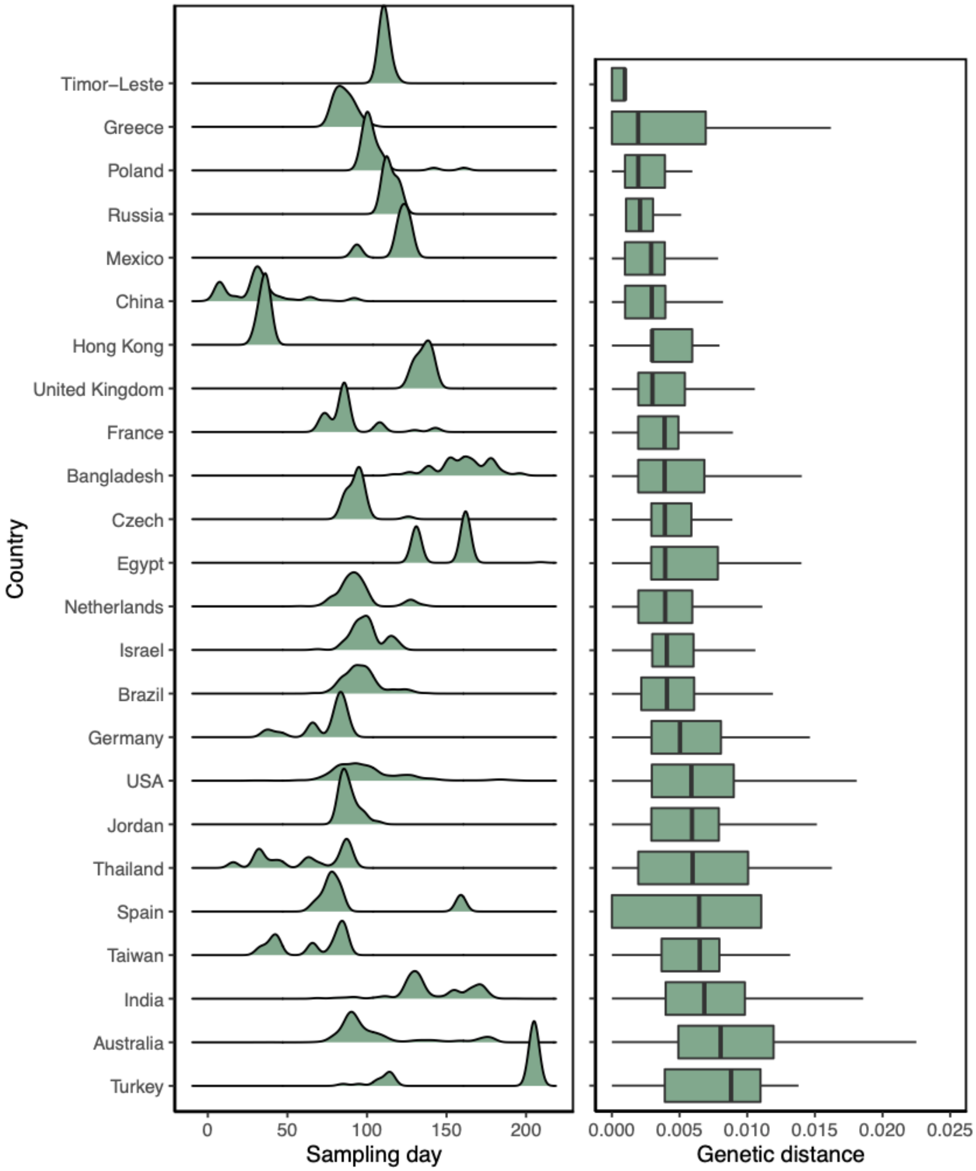
Country
Timor–Leste
Greece
Poland
Russia
Mexico
China
Hong Kong
United Kingdom
France
Bangladesh
Czech
Egypt
Netherlands
Israel
Brazil
Germany
USA
Jordan
Thailand
Spain
Taiwan
India
Australia
Turkey
0
50
100
150
200
Sampling day
0.000
0.005
0.010
0.015
0.020
0.025
Genetic distance
